# *Daphniphyllum macropodum* genome assembly and multiplexed screening unveils alkaloid modifying enzymes

**DOI:** 10.64898/2026.11.16.766977

**Authors:** Sandesh H. Swamidatta, Barbara A. Radzikowska, Danielle J. Taylor, Samuel J. Smit, Lachlan J. N. Waddell, Swen Langer, Tony R. Larson, William P. Unsworth, Benjamin R. Lichman

**Author notes:** Corresponding author Benjamin R. Lichman.

## Abstract

*Daphniphyllum macropodum*, a small shrub or tree native to East Asia, produces bioactive polycyclic alkaloids of remarkable complexity and variety, making them long-standing, attractive targets for chemical synthesis. However, the biosynthetic origins of *Daphniphyllum* alkaloids remain unknown, and no pathway-associated enzymes have been identified to date. Here, we report the chromosome-level genome assembly of *D. macropodum*, representing the first genome assembly for the Daphniphyllaceae, a family in the Saxifragales woody clade. To discover alkaloid-modifying enzymes, we developed a highly combinatorial, multiplexed screening approach which enabled testing multiple enzyme candidates with multiple substrates in a single reaction. Deconvolution and characterisation identified two cytochrome P450 enzymes that catalyse the hydroxylation of the yuzurimine-type alkaloid daphhimalenine B. Specifically, CYP82C230 functions as a daphhimalenine B 2-hydroxylase to yield the natural product yuzurimine C, whereas its paralog, CYP82C231, acts as a daphhimalenine B 13-hydroxylase generating a novel alkaloid we have named yuzurimine F. These are the first reports of an enzymatic modification of a *Daphniphyllum* alkaloid. Collectively, this work advances our understanding of *D. macropodum* biology and represents the first steps towards elucidating the complex *Daphniphyllum* alkaloid biosynthetic pathway.

## Introduction

Nature has a remarkable ability to forge complex molecular architectures from simple building blocks. These natural products can be used as pharmaceuticals, agrochemicals and consumer products. Resolving the chemical, genetic, and enzymatic pathways that form them provides us with new synthetic and biological tools for controlled fabrication of both natural products and related novel chemicals^1^.

The *Daphniphyllum* alkaloids (DAs) are highly complex terpene-alkaloids produced exclusively in plants from the genus *Daphniphyllum*^2^, notable for their three-dimensionality and stereochemical density (**Fig. 1**)^3–5^. Very little is known about how these compounds are formed in the plants. Identifying and characterising enzymes that contribute to the biosynthesis of DAs would reveal the strategies by which nature controls the formation and modification of complex structures, and provide us with new tools to make bioactive compounds.

**Figure 1.**
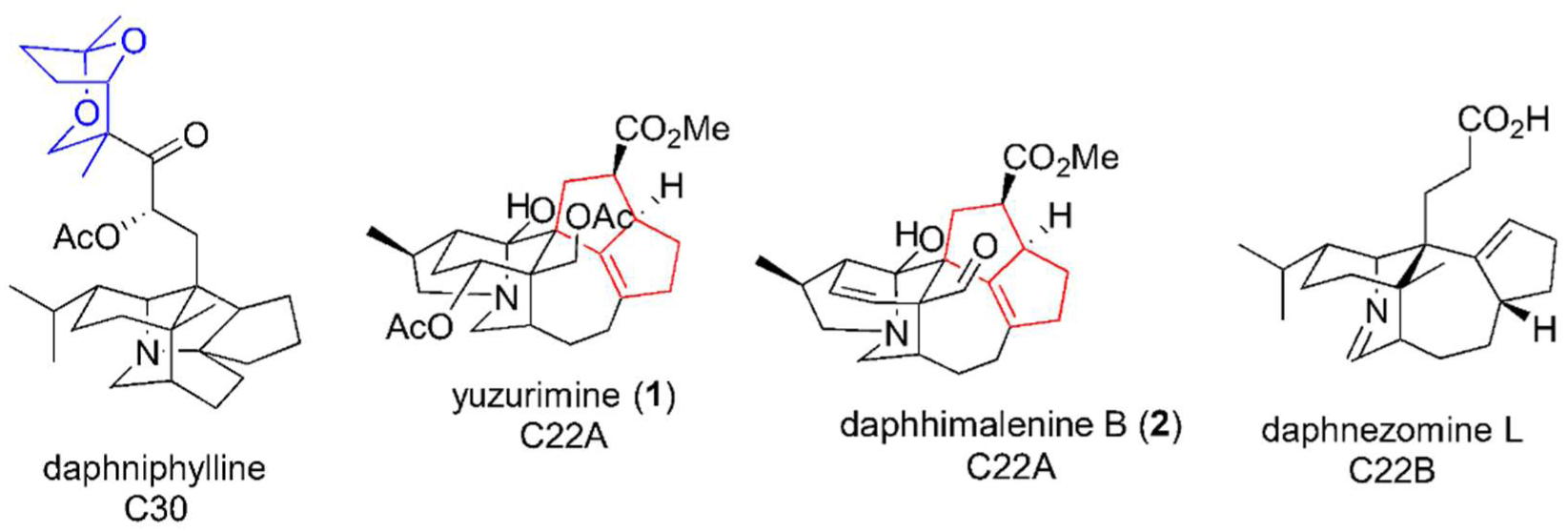
Representative *Daphniphyllum* alkaloids found in *D. macropodum.* Classified by carbon skeleton (C30 vs C22, blue highlighting C8 fragment lost in C22 compounds) and presence (C22A) or absence (C22B) of hexahydropentalene ring (red)^16^.

*Daphniphyllum* consists of around 30 species native to tropical and subtropical Asia^6^. It is the sole genus of the ancient Daphniphyllaceae family, which diverged from within the woody clade of the Saxifragales approximately 100 million years ago^7^. *Daphniphyllum macropodum* Miq., a dioecous evergreen tree or shrub, is the most widely known *Daphniphyllum* species, due to its worldwide cultivation for landscape horticulture. In Japan it is called Yuzuriha (ユズリハ), meaning “yielding leaves”, which refers to new leaves growing over the old prior to them falling off.

In China, some communities use *D. macropodum* leaves as treatment for skin conditions^8^, though the foliage is also known to be toxic to grazing cattle^9^. The roots of *Daphniphyllum calycinum* are used in Chinese medicine as a component of colitis treatment^10^. The therapeutic or toxic effects of plants can be attributed to their chemical composition, and accordingly the bioactivity of DAs have been investigated. Of the more than 350 isolated DAs, many have been shown to be bioactive^3,11^, including recent descriptions of antiviral activity^12^ and cancer cell cytotoxicity^13,14^.

The DAs can be classified into 35 subtypes based on their unique backbones including daphniphylline, secodaphniphylline, calyciphylline A and yuzurimine subtypes^15^. A broader classification based on the number of carbon atoms in the core skeleton groups DAs into C30, C22A and C22B types, with the C22s split based on the presence of a hexahydropentalene ring^16^ (**Fig. 1**). The polycyclic cage-like architectures of DAs have made them attractive targets for total synthesis^4^. Of particular note are Heathcock’s iconic biomimetic syntheses of the daphniphylline skeleton featuring a reaction cascade from a squalene-like precursor ^17,18^. The unique complexity of DAs ensures they remain popular targets in synthetic chemistry^19–21^.

Yuzurimine (**1**) (**Fig. 1**) is a major DA found in *D. macropodum*; its structure was first described in 1966^22^. Since then, many yuzurimine-type DAs have been isolated including yuzurimines A, B and C^23,24^, several of which have biological activities^25,26^. These yuzurimine-type compounds contain a complex tetracarbocyclic core framework and have been the subject of total synthesis campaigns^27^. Although significant advances in synthesis have enabled the construction of their complex cage-like cores in 21 steps^28^, total synthesis remains a challenge. Recently, the first total synthesis of a yuzurimine-type DA was reported: the asymmetric total synthesis of (+)-caldaphnidine^29^. Since then additional yuzurimine-type DA syntheses have been achieved including C14-epi-deoxycalyciphylline H, yuzurimine A and yunnandaphnine B^30,31^. Enzymes that can modify yuzurimine-type DAs would supplement the synthetic efforts exploring the structural and bioactivity potential of this chemical space.

The biological basis for DA formation and diversity remains a mystery. Most biosynthetic speculation has centred around the origin of the DAs from squalene, proposed by Suzuki^32^ and supported by Heathcock’s biomimetic syntheses^17,18^. Our recent work using metabolomics to investigate DA accumulation in *D. macropodum* supports the terpenoid origin of all DAs and furthermore revealed a distinct distribution patterns of C30 and C22 DAs between and within various tissues, suggesting that C30 DAs may be formed in the phloem prior to conversion to C22 DAs, such as yuzurimine, which accumulate in the epidermis^16^. We later characterised *D. macropodum* terpenoid-related enzymes but are yet to reveal the enzymatic link between terpenoid biosynthesis and DA formation^33^. Therefore, there are, as of yet, no reported enzymes capable of forming or modifying DAs.

Biosynthetic elucidation in non-model species and in atypical pathways is challenging, despite integrative omics-based approaches accelerating the success of pathway discovery in such plants^34^. Integration of genome sequencing with coexpression based approaches can greatly enhance identification of enzyme families, genomic features such as biosynthetic gene clusters and duplication events as well as revealing evolutionary contexts^35–37^. Furthermore, heterologous expression in *Nicotiana benthamiana* by expressing pools of genes enables rapid identification for multistep pathways^38,39^. This system also allows efficient combinatorial screening of enzymes by substrate feeding^40^.

Although there has been a huge interest in the chemical synthesis of DAs for decades, no biosynthetic enzyme in the pathway has so far been identified. To enable elucidation of the metabolic origin of DAs we first sequenced the genome of *Daphniphyllum macropodum* and generated a multi-omics datasets. Using these datasets, we are able to identify two cytochrome P450 enzymes capable of modifying the yuzurimine type scaffold. Our combinatorial “dual-pooling” of genes and metabolites for screening using *Nicotiana benthamiana* expression system combined with leaf disc based biocatalytic conversion resulted in biosynthesis and characterisation of a novel alkaloid. This study reports the first enzymatic modification of DA scaffolds and highlights how multi-omics approaches combined with combinatorial screening can produce new-to-nature compounds.

## Results

### Genome sequencing and assembly

We sequenced and assembled the genome of *Daphniphyllum macropodum* (2n = 32) using a combination of PacBio circular consensus sequencing (51.1 Gb reads; 55X coverage) and Dovetail proximity ligation “Omni-C” reads (30X coverage). The initial assembly, conducted with hifiasm^41^, was 1181.59 Mbp which is in agreement with the genome size of 1250 Mbp estimated through flow cytometry (**Supplementary Fig. S1**) and that of the related species *D. pentandrum*^42^. Phasing and scaffolding with Omni-C reads resulted in haplotype resolved assemblies of 1105.76 Mbp (haplotype 1) and 1114.35 Mbp (haplotype 2), with both assemblies anchored to 16 pseudomolecules (**Figure 2, Supplementary Fig. S2, Supplementary Table S1**). Both haplotypes had assembly BUSCO scores above 98% complete (**Supplementary Table S2**), indicating high quality chromosome-level assemblies. The respective BRAKER annotations using *ab initio*, homologous and transcription-based prediction provided annotations with BUSCO scores greater than 98% complete, with 96% single copy (**Supplementary Table S2**), encoding for 23,819 and 24,569 gene models, respectively. Repeats and transposable elements covered 74% of the genome, of which long terminal repeats (LTR) elements contributed 42% (**Supplementary Fig. S3, Supplementary Table S3**).

**Figure 2.**
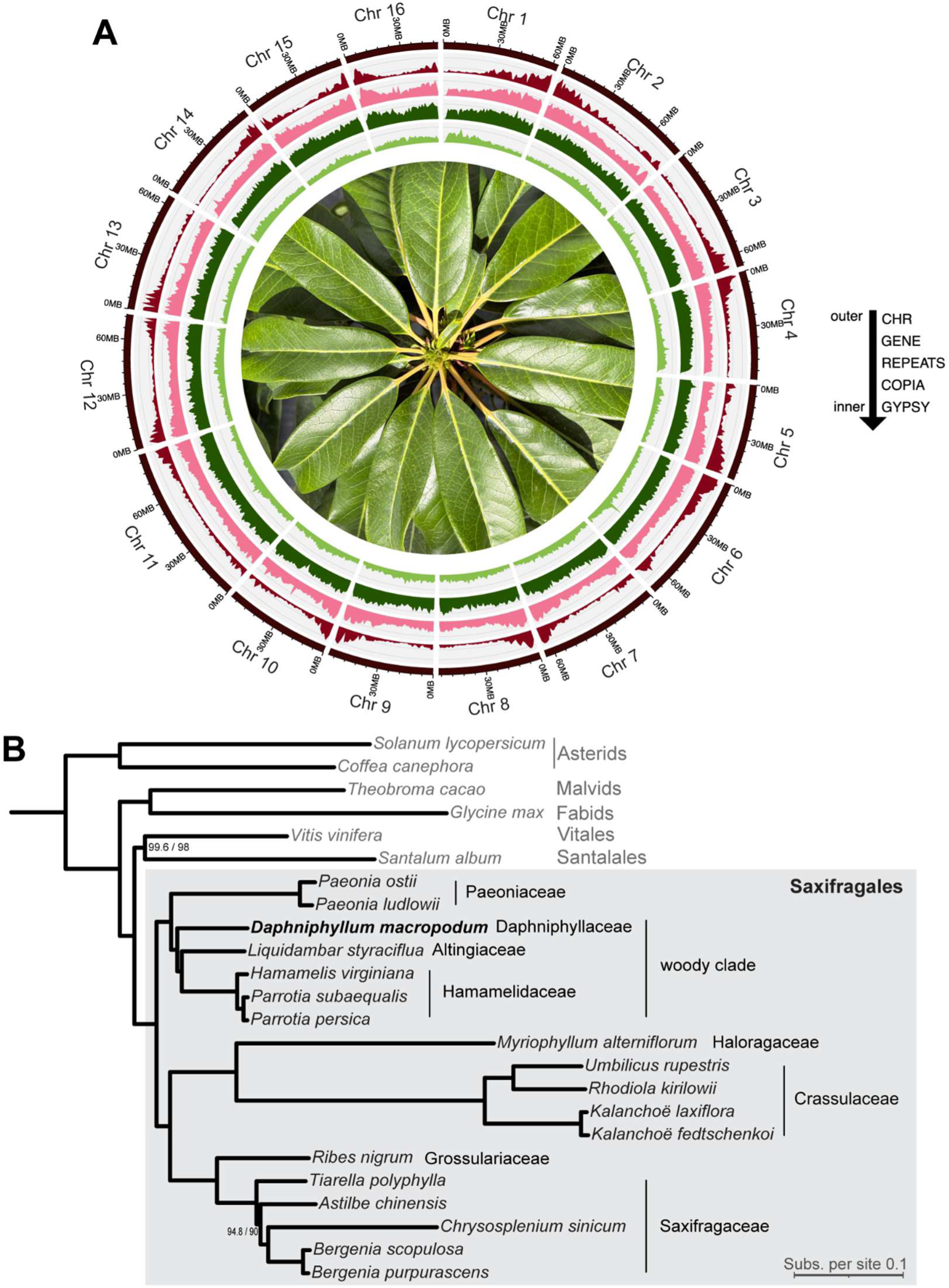
*Daphniphyllum macropodum* genome assembly and phylogenetic context. **A.** The genomic landscape of *Daphniphyllum macropodum* with 16 pseudochromosomes represented by circos plot. **B**. Maximum likelihood phylogeny of Saxifragales based on single copy nuclear genes from whole genome assemblies. The position of *Daphniphyllum macropodum* is shown in bold. Branch supports were calculated with ultrafast bootstraps / SH-aLRT, and are fully supported unless noted.

### Phylogenetic context of *Daphniphyllum macropodum*

This *Daphniphyllum macropodum* genome assembly is the first reported from the Daphniphyllaceae, so it provides a valuable resource for understanding the evolution of the diverse and ancient Saxifragales clade^43^. To examine the phylogenetic context of *Daphniphyllum macropodum*, we inferred a species-level maximum likelihood tree using whole genome assemblies, eighteen from Saxifragales and six outgroups representing major angiosperm clades (**Figure 2B**, **Supplementary Table S4**). The tree was rooted on core asterids based on the topologies recovered from recent major phylogenetic studies^44,45^. Saxifragales was recovered as a single clade, sister to a single clade containing Vitales and Santalales. Together this clade (Saxifragales, Vitales, Santalales) is sister to the core rosids (Malvids, Fabids). This topology contrasts with plastid derived phylogenies^46^ and a recent major phylogenomic study^45^, but discordance is not surprising as the relationships between superrosid taxa are notoriously challenging to resolve^44^ (**Supplementary Fig. S4A**). We also noted that the topology within Saxifragaceae was different from previous reconstructions, again highlighting the challenge of resolving relationships within this order (**Supplementary Fig. S4B**)^43,47,48^. Within the Saxifragales, Daphniphyllaceae is a member of the “woody clade” which includes the trees *Hamamelis virginiana* (witch-hazel) and *Liquidambar styraciflua* (sweetgum). Here, we resolve the woody-clade as sister to Paeoniaceae and Daphniphyllaceae as an early diverging member, which contrasts with plastid phylogenies^7,46^ but is in line with recent nuclear phylogeny studies (**Supplementary Fig. S4C**)^45,47^.

### Identification of *Daphniphyllum* alkaloid tailoring enzymes

Yuzurimine subtypes are one of the major subtypes of DAs with over 50 members^15^, including some with bioactivities^25,26^. We hypothesised that yuzurimine subtype diversity arises from tailoring steps involving site-selective hydroxylations and subsequent acetylation reactions. Such selective tailoring reactions are challenging to perform synthetically, however in plant biosynthesis they can be achieved through the action of oxidative enzymatic strategies (such as cytochrome P450 catalysed reactions) which, in turn, can facilitate biocatalytic routes for chemoenzymatic synthesis^49^. Therefore we sought to identify enzymes that can modify yuzurimine-type scaffolds. In our previous work, C22A DAs, which includes yuzurimine subtypes, were found to show differential distribution and enrichment in aerial organs (mainly in immature leaves and petioles) with tissue localisation in the epidermis^16^.

To exploit differential metabolite accumulation for gene discovery, we generated paired metabolomics and transcriptomic datasets for *D. macropodum* across six tissue types (bark, immature leaf, mature leaf, petiole, stem and root) (**Supplementary Table S5**). The transcriptomic data was mapped to the genome gene models, and in the metabolomic data the peak representing yuzurimine (**1**) was identified through an NMR verified standard (**Fig. 3A**). We performed a Spearman’s correlation analysis to correlate the accumulation of yuzurimine with transcriptome data set to identify potential candidate genes capable of tailoring yuzurimine subtype DAs (**Fig. 3B**). The list was filtered for correlation with yuzurimine (*r* > 0.6), highest expression in immature leaf and petiole, and for PFAM annotations corresponding to tailoring enzymes (cytochrome P450, 2OG-Fe(II) oxygenase, transferase and dioxygenase). In addition, we also identified a candidate (*Dm_H1_g4974.t1*) showing high correlation with the top yuzurimine-correlating candidate, *Dm_H1_g8422.t1*, resulting in a set of 16 candidate genes (**Fig. 3B**, **Supplementary Table S6**).

**Fig. 3.**
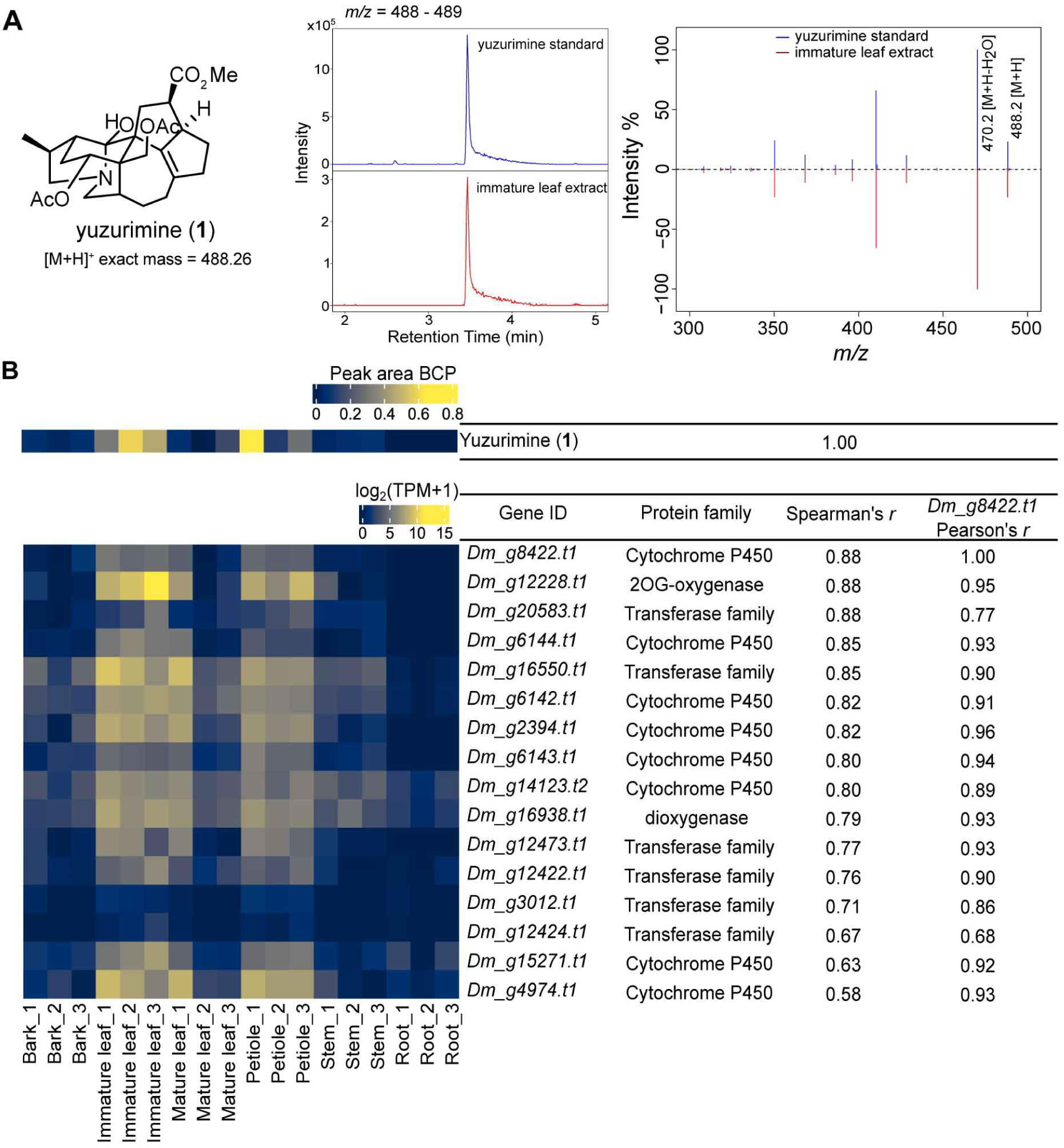
Metabolite to gene correlation analysis for yuzurimine. **A.** Yuzurimine (**1**) structure, extracted ion chromatogram (*m/z* 488-489), and MS^2^ mirror plot confirming yuzurimine (**1**) identity in samples. **B.** Heatmap of selected candidate tailoring enzymes with expression patterns similar to accumulation of yuzurimine as identified using Spearman’s correlation (*r*). A gene-to-gene correlation (Pearson’s *r*) of selected candidate genes with top hit (*Dm_g8422.t1*) from metabolite-gene correlation is also shown. BCP = blank corrected percentage, TPM = transcripts per million, r = correlation coefficient.

### A dual-pooled screening approach combined with untargeted metabolomics identifies a DA tailoring Cytochrome P450

To screen candidate enzymes for the ability to modify DAs, we employed a “dual-pooled screening” approach which leverages the power of pooling genes and pooling substrates simultaneously in a single assay (**Fig. 4**). To pool genes, we used the powerful *Nicotiana benthamiana* transient gene expression system which allows for mixing multiple *Agrobacterium* cultures each containing a unique gene expression vector into a single infiltration, leading to a single leaf expressing multiple gene candidates^39^. After four days, leaf discs were excised and incubated with *D. macropodum* alkaloid extracts containing mixtures of alkaloids^40^. This approach conserves stocks of purified compounds which may have limited availability and also allows the enzymes to act on the pool of potential substrates present in the mixed extracts. After incubation, the discs and their media are analysed by untargetted metabolomics using high-resolution LC-MS, and the data processed through software (XCMS) which identifies peaks with significant changes (i.e. decrease or increase) during incubation^50^.

**Fig. 4.**
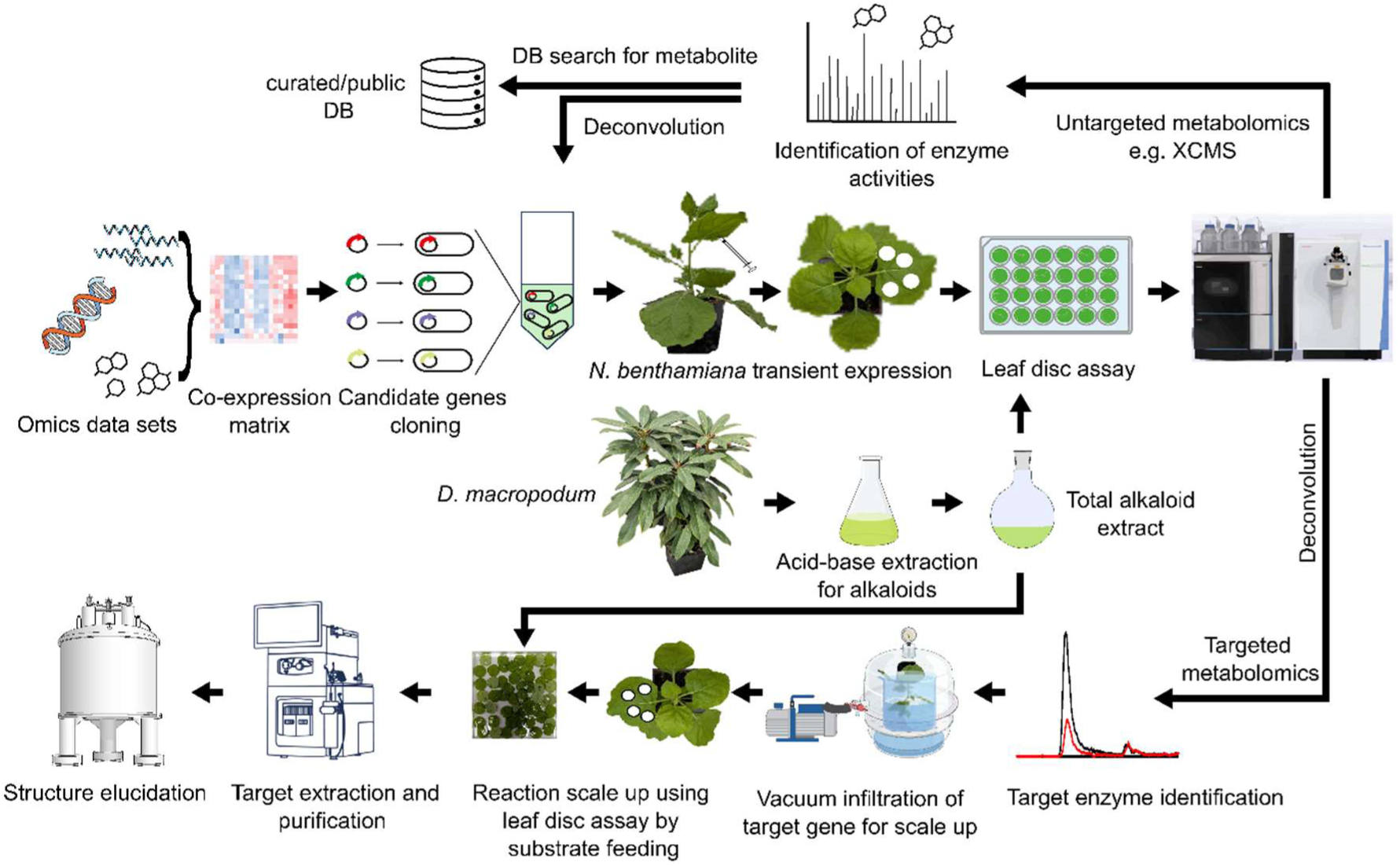
Dual-pooled multiplexed screening workflow.

We used the “dual-pooled” approach to screen two gene candidate pools: the first, a pool containing the 16 candidate genes identified through correlations (see **Fig. 3**) as well as two genes (*Dm_H1_g6141.t1* and *Dm_H1_g6140.t1*) derived from a cytochrome P450 array (**Fig. 5A**, **Supplementary Table S6**). The metabolomics analysis identified a peak with *m/z* 384.21 showing significant decrease compared to an empty vector control, suggesting this compound was a substrate for one or more of pooled enzymes (**Fig. 5B**). We also identified a peak with mass *m/z* 400.21 showing a significant increase compared to empty vector control suggesting product formation (**Fig. 5C**). The mass difference of these two peaks is 16, corresponding to an oxygen atom and indicating that the conversion is likely a hydroxylation. We then examined pool 2, a subset of pool 1 that contained only oxygenase genes (**Fig. 5A**). The same pattern of significant peak changes was observed (**Fig. 5D** and **5E**), suggesting that the causal gene/s involved must be in pool 2. To identify the active genes we then employed a gene drop-out strategy using multiple pools derived from the pool 2 set, with each pool generated by systematically omitting one gene at a time. The leaf disc assay was repeated on each drop-out pool by feeding total alkaloid extract from leaf tissue. Each pool showed activity except when *Dm_H1_g15271.t1* was omitted (**Fig. 5F** and **5G**, **Supplementary Fig. S5**) revealing that the cytochrome P450 encoded by *Dm_H1_g15271.t1* was the active enzyme. This enzyme was given the name CYP82C230.

**Fig. 5.**
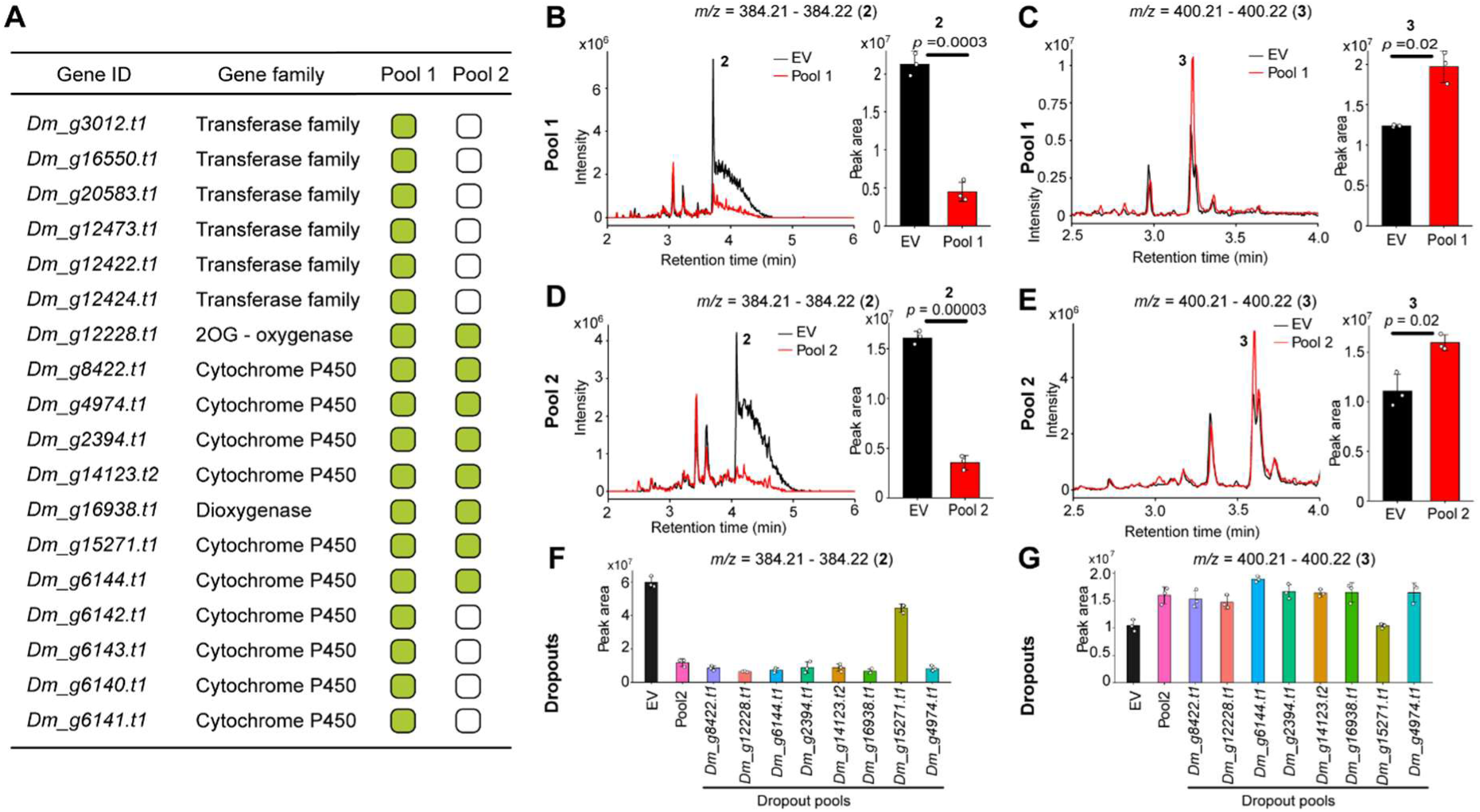
Identification of enzyme activities by ‘dual-pooled’ screening. **A.** List of candidate genes screened as two pools. The green boxes indicate the inclusion of the gene in that pool. **B-E.** Identification of peak changes in pooled screens. **B** and **C**. Pool 1 screen. **D** and **E**. Pool 2 screen. **B** and **D.** Reduction in peak *m/z* 384.2. **C** and **E.** Increase in peak *m/z* 400.2. **F** and **G**. Dropout test showing proposed substrate peak *m/z* 384.2 and product peak *m/z* 400.2 respectively. The bar charts show the changes in peak area for each mass feature between empty vector and pooled samples. The values represent mean ± SD (*n* = 3), two-tailed Student’s *t*-test.

### CYP82C230 is a daphhimalenine B 2-hydroxylase

Having identified the active gene and the likely substrate-product pair using the dual-pooled screening method and gene drop-out approach, we then characterised the enzyme activity through identifying the substrate and product. We were able to identify the apparent substrate peak (with *m/z* 384.21) was daphhimalenine B (**2**), based on an NMR verified standard (**Supplementary Fig. S6-S13**). To confirm the activity of CYP82C230 on daphhimalenine B (**2**), we repeated the leaf disc assay by expressing *Dm_H1_g15271.t1* / CYP82C230 on its own and tested its activity by feeding purified daphhimalenine B (**2**) (**Fig. 6**). The LC-MS/MS analysis confirmed the new peak with *m/z* = 400.21 was yuzurimine C (**3**), matching an NMR-verified standard purified from *D. macropodum* leaf tissue (**Fig. 6A-C**, **Supplementary Figs. S14-S19**). The structure of yuzurimine C (**3**) was previously elucidated by Hirata and co-workers, but only its ^13^C NMR data and selected ^1^H NMR signals were reported^24^. We have now completed a more comprehensive analysis and assigned all ^13^C and ^1^H signals using 2D NMR experiments (COSY, HSQC, HMBC) (**Supplementary Figs. S14-S19**). We further confirmed CYP82C230 enzyme activity through whole-cell feeding of daphhimalenine B (**2**) to a *Saccharomyces cerevisiae* strain expressing *Dm_H1_g15271.t1* / CYP82C230 (**Fig. 6A** and **D**). This characterisation of CYP82C230 as daphhimalenine B 2-hydroxylase represents the first demonstration of an enzymatic modification of a *Daphniphyllum* alkaloid.

**Fig. 6.**
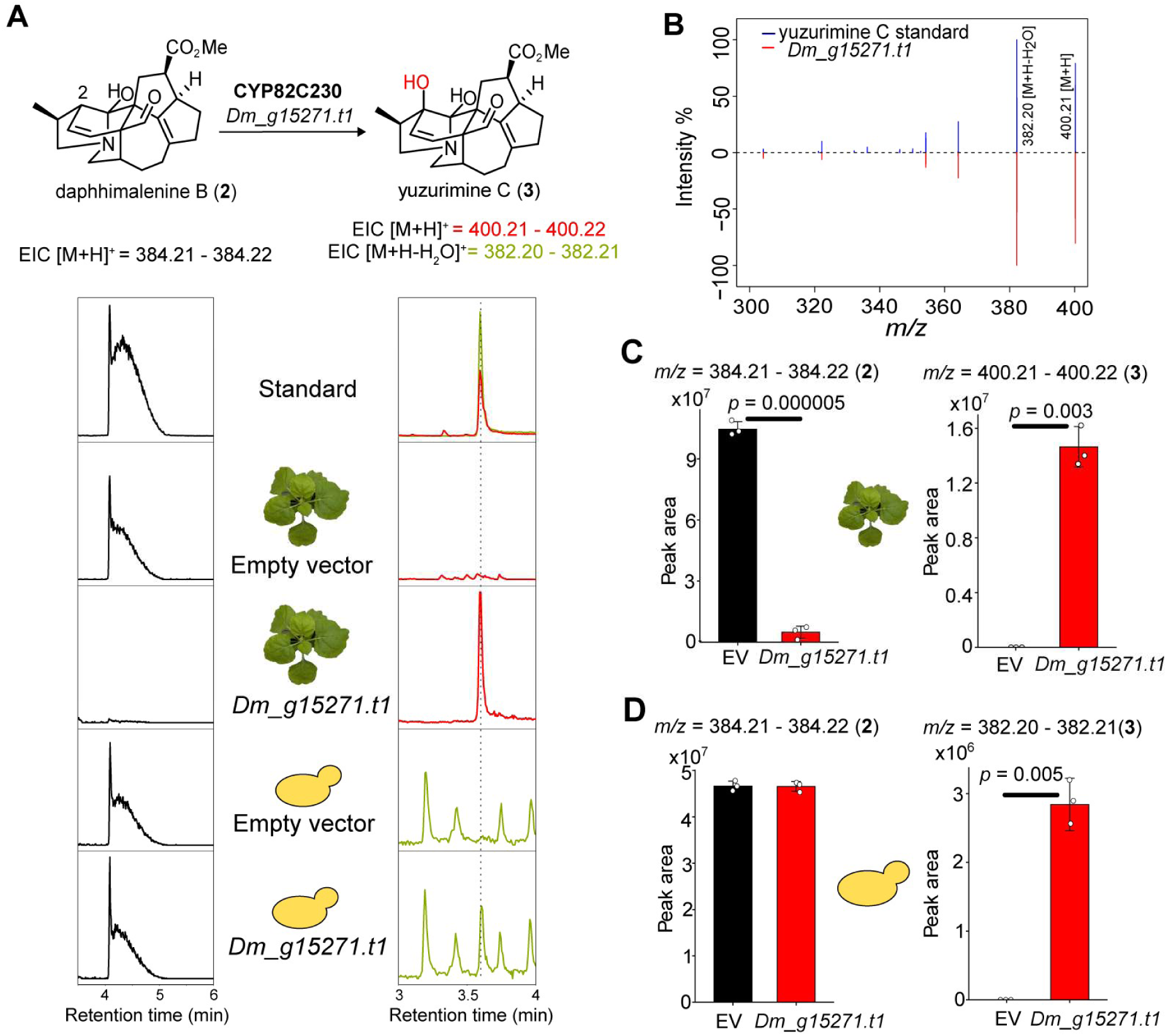
Confirmation CYP82C230 is a daphhimalenine B 2-hydroxylase. **A.** LCMS chromatograms showing conversion of daphhimalenine B (**2**) (*m/z* EIC 384.21 [M+H]^+^) to yuzurimine C (**3**) (*m/z* EIC 400.21 [M+H]^+^ and 382.20 [M+H-H_2_O]^+^) in the presence of the *Dm_H1_g15271.t1* gene in both plant and yeast chassis. **B.** MS^2^ mirror plot matching *N. benthamiana* produced yuzurimine C (**3**) with the NMR verified standard. **C** and **D.** Bar charts represent the changes in peak areas of daphhimalenine B (**2**) and yuzurimine C (**3**) between empty vector and *Dm_H1_g15271.t1* in *N. benthamiana* (**C**) and yeast (**D**). The values represent mean ± SD (*n* = 3), two-tailed Student’s *t*-test.

### A cytochrome P450 gene array reveals CYP82C231

Genes involved in plant specialised metabolism are often coexpressed and display a continuum of genomic organisations including tandem arrays, gene pairs, and biosynthetic gene clusters^37,51^. With this mind, we explored the genomic context of *Dm_H1_g15271.t1* and its association with potential biosynthetic genes using our new *D. macropodum* genome assembly. This revealed that the *Dm_H1_g15271.t1* is part of a six gene cytochrome P450 array, all paralogues showing over 70% pairwise protein sequence identity (**Fig. 7A**). Gene arrays, sets of homologous genes in close physical proximity on the genome, are a common feature of plant specialized metabolism^37,52,53^. These are typically generated by duplication events and facilitate expansion and diversification of enzymatic functions within biosynthetic pathways^54–56^. Hence, we hypothesised that the other array genes may show activities on DAs. To identify any functionalities, we repeated the leaf disc assay by individually expressing each cytochrome P450 and feeding leaf alkaloid extracts. This resulted in identifying an activity for *Dm_H1_g15275.t1* in which daphhimalenine B (**2**) appeared to be consumed (**Fig. 7B** and **7C**) and a new peak emerged with *m/z* 400.21, with the mass difference of 16 again indicating hydroxylation activity (**Fig. 7D** and **7E**). The new peak was different to yuzurimine C (**3**) and had not been observed in any of our metabolomics data sets. The enzyme encoded by *Dm_H1_g15275.t1* was named CYP82C231.

**Fig. 7.**
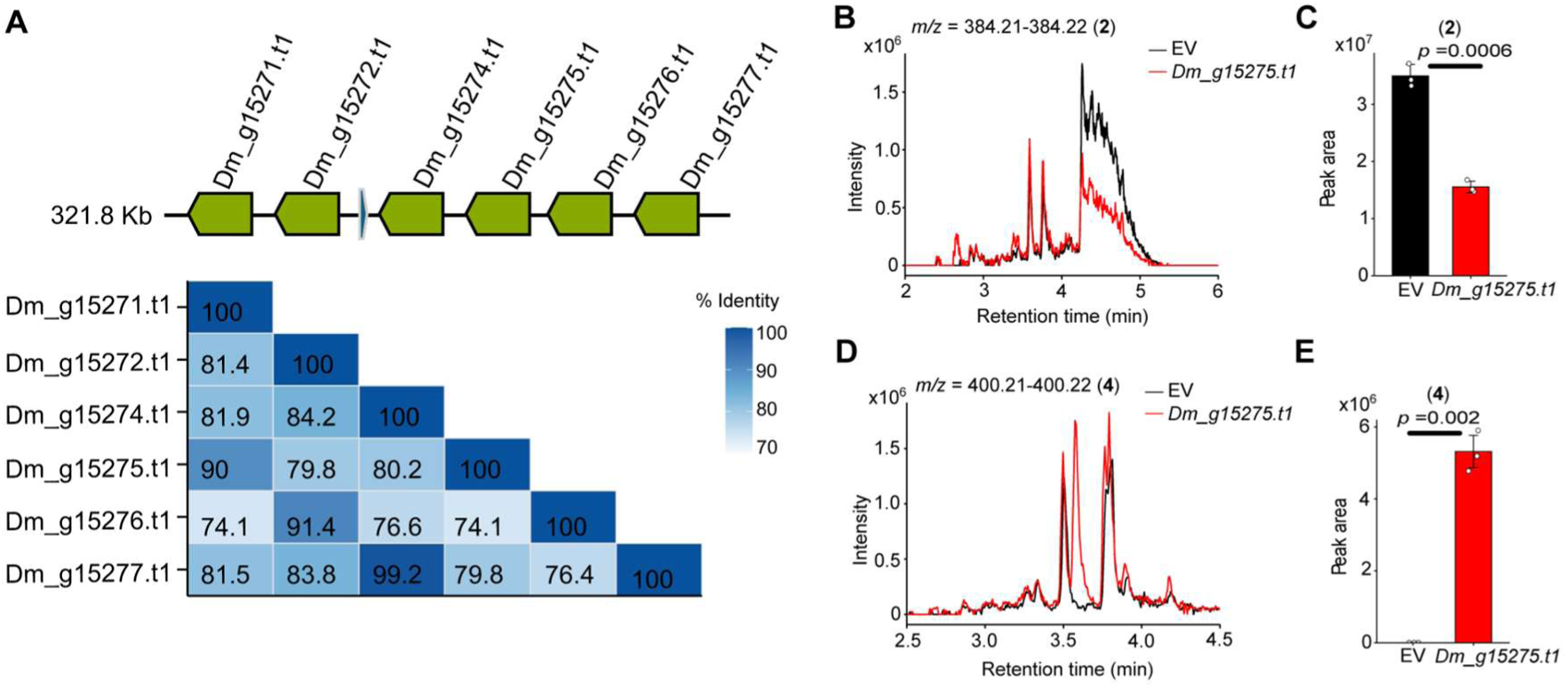
Screening the cytochrome P450 array. **A.** Schematic showing the cytochrome P450 array along with protein identity matrix of all six array genes. **B**-**E**. Identification of *Dm_H1_g15275.t1 /* CYP82C231 activity showing the consumption of a mass feature *m/z* = 384.21 (**B** and **C**) with appearance of a new mass feature m/z=400.21 (**D** and **E**). The bar charts (c and e) represent the mean peak areas of mass features. The values represent mean ± SD (*n* = 3), two-tailed Student’s *t*-test.

### CYP82C231 is a daphhimalenine B 13-hydroxylase

To isolate the new product we scaled-up the leaf disc assay, using it as a living whole-cell biocatalytic system (**Fig. 4**) ^57^. This approach ensures more efficient substrate utilisation compared to conventional infiltration of substrates into *N. benthamiana* leaf. For scaling up the reaction, harvested leaf discs expressing *Dm_H1_g15275.t1 /* CYP82C231 were fed with a semi-pure daphhimalenine B (**2**) (∼ 4 mg). After incubation, the product was extracted and purified; we were able to isolate 1.8 mg of product **4**. The structure was elucidated by 1D and 2D NMR (**Supplementary Fig. S20-25**). Initial analysis of ^1^H, ^13^C and HSQC spectra showed that a tertiary carbon of daphhimalenine B (**2**) had been replaced by a quaternary carbon. Thus, we deduced that hydroxylation had occurred at either the C-15 or C-6 positions. Further analysis of the COSY spectrum revealed that H-14 only showed *J* couplings with H-13a/b, while H-16a/b coupled with H17a/b. Therefore, we inferred that hydroxylation had occurred at the C-15 position. All expected H-H correlations for this proposed structure were observed in the COSY spectrum (**Supplementary Fig. S23**), while the HMBC spectrum provided further evidence by revealing numerous multi-bond C-H correlations consistent with our assignment (**Supplementary Fig. S24**). To the best of our knowledge this compound has not been isolated previously and represents a novel alkaloid, which we have named compound yuzurimine F (**4**). Using daphhimalenine B (**2**) as the substrate, and yuzurimine F (**4**) as the product standard, we were able to confirm the activity of CYP82C231 using both *N. benthamiana* leaf disc assays and yeast whole cell conversion (**Fig. 8**). CYP82C231 was confirmed to be a daphhimalenine B 13-hydroxylase.

**Fig. 8.**
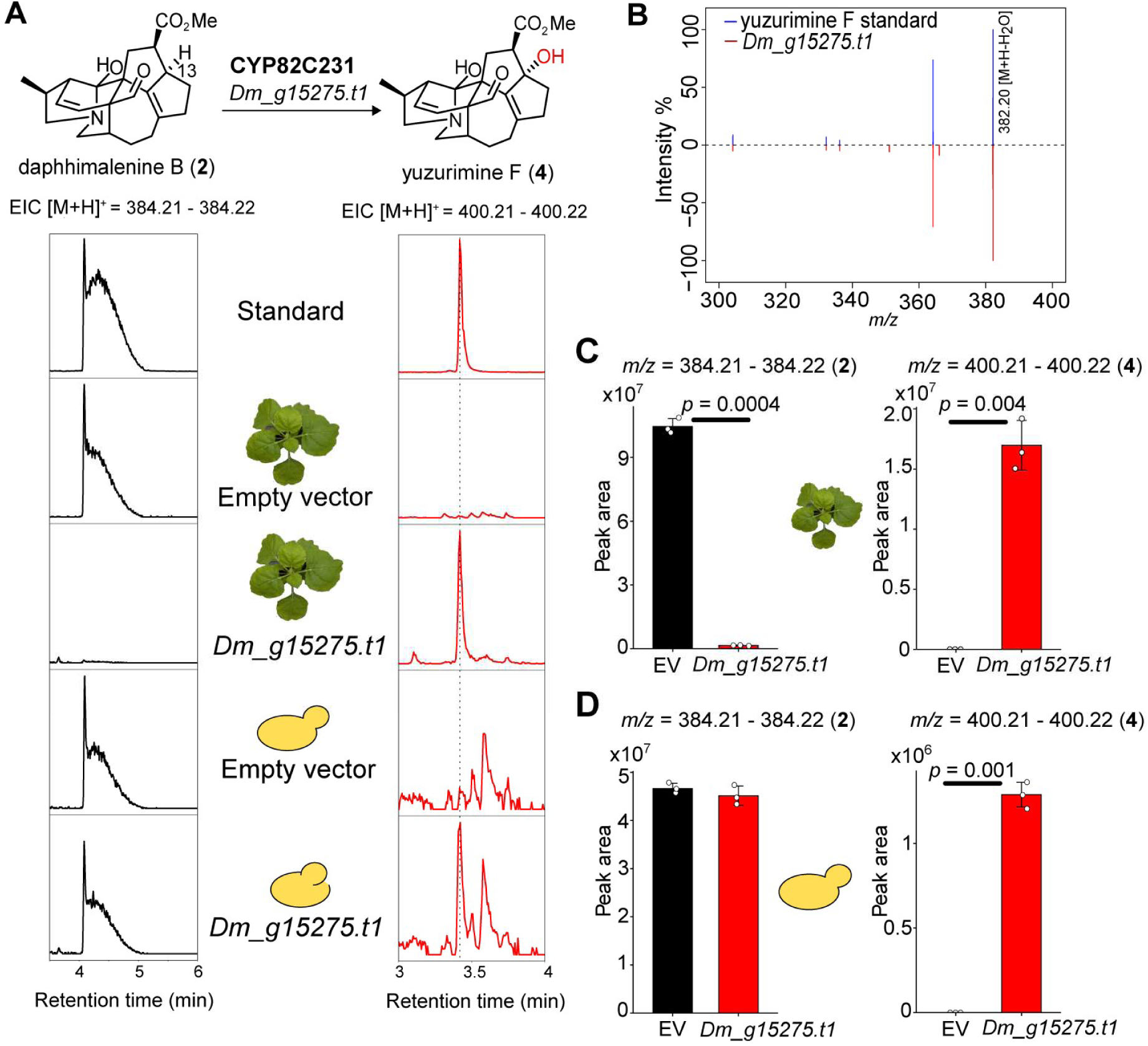
Confirmation CYP82C231 is a daphhimalenine B 13-hydroxylase. **A.** LCMS chromatograms showing conversion of daphhimalenine B (m/z EIC 384.21 [M+H]) to yuzurimine F (m/z EIC 400.21 [M+H]^+^) in the presence of *Dm_H1_g15275.t1* in both plant and yeast background. **B.** MS^2^ mirror plot matching yuzurimine F with the standard. **C** and **D**. Bar charts represent the changes in peak areas of daphhimalenine B (**2**) and yuzurimine F (**4**) between empty vector and *Dm_H1_g15275.t1* in *N. benthamiana* (**C**) and yeast (**D**). The values represent mean ± SD (*n* = 3), two-tailed Student’s *t*-test.

## Discussion

The genomes of plants that produce complex bioactive compounds contain genes that encode a wealth of enzymatic machinery. This includes specific genes that control conversion of a simple precursor to a complex target product; discovering these pathway genes enables the reformulation of pathways in heterologous organisms leading to controlled production of a high-value product^58^. The genomes also contain genes that contribute more broadly to chemical diversity, notably through gene expansions^59^ and gene recruitment^60^.

Here, we investigated *Daphniphyllum macropodum*, a plant that produces highly complex and unique squalene derived terpene-alkaloids, obtaining a genome assembly in order to access enzymatic machinery that could aid in the exploration of a largely uncharted chemical space. This interdisciplinary work begins with a chromosome-level genome assembly of a non-model plant, and leads to the biocatalytic formation and isolation of a novel alkaloid, via a workflow leveraging the combinatorial power of *N. benthamiana* and untargetted metabolomics.

We report the first chromosome-level assembly for a species in the *Daphniphyllum* genus, and, by extension, the first assembly for a species in Daphniphyllaceae. This has implications beyond natural product research. The assembly from an ancient lineage (diverging from within the Saxifragales woody clade approximately 100 million years ago^7^) will aid in the understanding of the evolutionary relations within and close to Saxifragales^43,47^, an order that originated around 150 million years ago, when multiple angiosperm lineages diverged during an event referred to as “Darwin’s abominable mystery” ^45^. This genome may also aid studies looking into the adaptation of *Daphniphyllum* across different latitudes^61^ and altitudes^62^, now allowing a genetic basis of adaptation to be investigated.

Effective and efficient discovery of genes encoding enzymes able to perform desirable chemical transformations requires consideration of multiple factors including candidate selection, throughput and sensitivity. In this work we were able to discover new oxidative activities on DAs by selecting gene candidates through gene-metabolite correlation, functional annotation, and array membership. After candidate selection, we were able to increase the effective throughput of the screening experiment by using a dualed pool approach, grouping both gene and substrate candidates in single experiments and subsequently deconvoluting them using drop-out or metabolomics analysis respectively. The *Nicotiana benthamiana* transient expression system is a highly versatile tool for combinatorial biosynthesis investigations, enabling simultaneous multiple gene infiltrations for pathway reconstruction and/or gene candidate screening^63^. Here we leveraged recent advances using leaf discs for feeding substrates^40^ and performing biotransformations^57^, We also extended these methods by feeding mixed alkaloid extracts and using metabolomics analysis to identify conversions in these mixtures, thereby enabling multiple genes and multiple substrates to be explored in a single sample. This approach effectively increases the throughput of the experimental system and allows unbiased discovery of new enzyme reactions.

We used this method to identify two novel cytochrome P450s: daphhimalenine B 2-hydroxylase (CYP82C230) and daphhimalenine B 13-hydroxylase (CYP82C231). These are the first reported enzymes capable of modifying DAs. The enzymes are from the CYP82 family, which is unique to angiosperms and is part of the large 71-clan^64^. CYP82 are commonly associated with specialised metabolism including those involved in benzylisoquinoline alkaloid biosynthesis^65^ and CYP82C2, a pathogen-induced enzymes responsible for the formation of the phytoalexins in Arabidopsis ^66^. CYP82C230 catalyses the hydroxylation of daphhimalenine B (**2**) to yuzurimine C (**3**) and may be the enzyme responsible for this step in the plant.

CYP82C231 catalyses the hydroxylation of daphhimalenine B (**1**) to yuzurimine F (**4**), which is not a previously reported natural product but a “new-to-nature natural product”^67^. The ability to make “new-to-nature natural products” using genes derived from a single genome highlights that its biosynthetic potential is not solely apparent through metabolite content: there may be many potential conversions which can only be revealed by functional screening. Furthermore, observing conversions is a peek behind the curtain into an “underground metabolism”^68^ where gene expansions of active enzymes can lead to the generation of enzyme variants with modified activities^69^. This process can fuel metabolic diversity^70^.

Heathcock’s formation of a *Daphniphyllum* alkaloid (DA) skeleton through a cascade reaction of an oxidised squalene derivative is an iconic demonstration of biomimetic molecular manipulation^17^. Whilst it is clear that the DAs derive from squalene^32^, we are yet to elucidate the specific biological or enzymatic processes responsible for this. Whilst the genome assembly and associated tissue atlas will be crucial for this work going forward, it has not yet provided an immediate solution to the origins of DA biosynthesis. This is due to multiple challenges including the inability to identify the biosynthetic precursors *in planta*, the lack of clear “bait” gene to inform co-expression analysis or gene cluster identification, and the ubiquity of DAs across all tissues^16^. The observation that DAs demonstrate within-organ distribution points suggests that single-cell transcriptomics may be the most powerful tool for unlocking this treasure chest of natural products^71,72^.

## Methods

### Plant materials and growth conditions

*Nicotiana benthamiana* plants were grown on F2 + S seed and modular compost (Levingston Advance) and kept and maintained for 4 to 5 weeks in the growth room. The growing conditions were 16-hour photoperiod and a temperature of 22 °C during the day and 20 °C during the night. For the genome sequence plant, *Daphniphyllum macropodum* Miq. seeds were collected from Castle Howard, North Yorkshire, UK (accession 19879448, plant B, planted 1989) and germinated and grown hydroponically as previously described^16^. Plants were transferred to pots with F2 + S seed and modular compost to establish. The offspring of plant B (labelled “B3”) was used for genome sequencing and the accompanying tissue atlas RNAseq for gene annotation. For paired transcriptome-metabolome analysis, tissues were obtained from three potted plants originally obtained from Architectural Plants, West Sussex, UK and maintained at the University of York (plants Fa, Fb and Fc, planted 2018)^16^.

### RNA extraction and sequencing

Total RNA was extracted from ∼100 mg frozen, finely ground plant tissue using a modified Cetyltrimethylammonium bromide (CTAB) method followed by silica column purification. Preheated (65 °C) extraction buffer (2% (w/v) CTAB, 100 mM Tris–HCL (pH 8), 2 M sodium chloride (NaCl), 2% (w/v) polyvinylpyrrolidone, supplemented immediately with 2% β-mercaptoethanol and 0.0025% Spermidine trihydrochloride before use) was added to ground tissue, vortexed thoroughly, and incubated at 65 °C for 10 min with intermittent mixing. Samples were cooled briefly and extracted twice with chloroform:isoamyl alcohol (24:1), followed by centrifugation at 12,000 ×g for 20 min at room temperature. The aqueous phase was recovered carefully after each extraction. RNA was precipitated by addition of 0.25 volumes 10 M LiCl and incubated at 4 °C overnight. Then RNA was pelleted (14,000 ×g, 40 min, 4 °C), washed with ice-cold absolute ethanol, and resuspended for purification using the RNeasy Plant Mini Kit (Qiagen) according to the manufacturer’s instructions. To remove residual contaminants and genomic DNA, samples were further purified using the RNA Clean & Concentrator-5 kit (Zymo Research) with on-column DNase I digestion, following the manufacturer’s protocol. Finally, RNA was eluted in 25 µL nuclease-free water and RNA quantity and quality were assessed using the Agilent Tapestation system with RNA nano screentape.

For short read sequencing, library preparation was performed from 200 ng high quality total RNA input using the NEBNext Ultra II Directional Library prep kit for Illumina in conjunction with the NEBNext Poly(A) mRNA Magnetic Isolation Module and unique dual index set (New England Biolabs, Ipswich, MA, USA), according to the manufacturer’s instructions. Libraries were pooled at equimolar ratios and sent for paired end 150 base sequencing at Novogene (UK) Company Ltd. on a NovaSeq 6000 instrument (Illumina, San Diego, CA, USA), with 10 million reads per sample on average.

RNAseq reads were quantified using SALMON v0.14.2. The transcript index was created using the whole genome as decoy followed by quantification of reads using automatic library inference and quasi-mapping validation (--validateMappings flag). Transcripts per million (TPM) for each mapped library were aggregated to generate an expression matrix.

### Genome sequencing and Assembly

The genome size and ploidy was estimated by flow cytometry (FCM) using TrisMgCl2 buffer according to the method of ^73^. A CytoFLEX LX (Beckman Coulter) FCM with a 561 nm excitation laser, 610/20 emission filter, and a flow rate of 30 μL/min was used. The threshold was set to 488 nm forward scatter to exclude instrument noise and background signal from the buffer. Fluorescence intensities of stained nuclei were used to analyze the nuclear DNA content and determine the genome size relative to the reference samples of *Glycine max* polanka (2C = 2445 Mbp) and *Zea mays* CE-777 (2C = 5311 Mbp).

Young leaves were ground under liquid nitrogen followed by extraction of high molecular weight DNA from 1 g of tissue using the NucleoBond HMW DNA Extraction kit (Macherey-Nagel, Düren, Germany). The DNA quality was assessed using Agilent Tapestation (running genomic DNA screentape). DNA samples were quantified using a Qubit 2.0 Fluorometer (Life Technologies, Carlsbad, CA, USA). The PacBio SMRTbell library (∼20 kb) for PacBio Sequel was constructed using SMRTbell Express Template Prep Kit 2.0 (PacBio, Menlo Park, CA, USA) using the manufacturer recommended protocol. The library was bound to polymerase using the Sequel II Binding Kit 2.0 (PacBio) and loaded onto PacBio Sequel II. Sequencing was performed on PacBio Sequel II 8M SMRT cells.

Nuclear DNA was extracted from leaf tissue and fixed in place with formaldehyde followed by Dovetail Omni-C library preparation. Fixed chromatin was digested with DNase I and then extracted, chromatin ends were repaired and ligated to a biotinylated bridge adapter followed by proximity ligation of adapter containing ends. After proximity ligation, crosslinks were reversed and the DNA purified. Purified DNA was treated to remove biotin that was not internal to ligated fragments. Sequencing libraries were generated using NEBNext Ultra enzymes and Illumina-compatible adapters. Biotin-containing fragments were isolated using streptavidin beads before PCR enrichment of each library. The library was sequenced on an Illumina HiSeqX platform to produce ∼ 30x sequence coverage.

PacBio CCS reads were assembled with HiC-Hifiasm using default parameters. Contaminant contigs were removed from the Hifiasm assembly using blobtools2 v1.1.1. Finally, purge_dups3 v1.2.5 was used to remove haplotigs and contig overlaps resulting in two haplotype assemblies. Omni-C reads were aligned to the respective haplotype assemblies using bwa (https://github.com/lh3/bwa) followed by scaffolding with HiRise ^74^ to generate the pseudomolecule haplotype resolved assemblies. Contiguity and completeness of the assemblies were assessed with BUSCO (Benchmarking for University Single Copy Orthologs) v5.4.2 with the embryophy-ta_odb10 dataset^75^.

### Genome Annotation

Genomic repeats and transposable elements (TEs) were masked with RepeatMasker v4.1.4 (using flags -nolow, -no_is and -xsmall) using a *de novo* library prepared with RepeatModler v2.0.4, including the LTR discovery pipeline. The masked genome was annotated with BRAKER3 using protein and RNA-seq data^76–85^. Viridiplantae proteins from OrthoDB v11 were used for protein evidence^86^. RNA-seq reads from bark, bud, immature leaf, mature leaf, petiole, root, stem and woody stem tissues were mapped to the masked genome using HiSAT2 v2.2.1 and converted to a BAM format using SAMtools v.1.20^87,88^. The resulting annotation was standardised and the longest transcript per gene model identified using AGAT^89^. The annotation quality was assessed using BUSCO v5.4.2. Transcripts were quantified as transcripts per million (TPM) by mapping RNA-seq reads using SALMON v0.14.^90^. Translated transcripts were annotated for functional domains using Interproscan v5.71-102.0^91,92^. Earl Grey v6.3.1^93^ using the Dfam v3.9^94^ transposable element database was used to comprehensively annotate and analyse repeats and TEs for the respective haplotypes. Gene, repeat and TE densities were determined with BEDtools v.2.30.0^95^ using 1Mb bins and plotted using shinyCircos^96^.

### Species phylogeny of Saxifragales

Genome assemblies for eighteen Saxifrigales, including *D. macropodum*, and six outgroup species (**Table S4**) were used to identify single-copy nuclear (SCN) genes. GFF annotations for the respective assemblies were used to extract primary coding sequences with the AGAT tool^89^. SCN genes were identified with MarkerMiner v.1.0^97^ using *G. max* as the single copy reference. The resulting MAFFT^98^ alignments where at least twenty-three of twenty-four species are present were trimmed for gaps using the gappyout algorithm of trimAl^99^. The trimmed alignments were concatenated into a supermatrix with partitions using the catfasta2phyml script (https://github.com/nylander/catfasta2phyml). IQ-TREE 2 was used to infer a maximum likelihood species tree with ModelFinder, ultrafast bootstraps (UFBoot2, X1000) and SH-aLRT supports (X1000)^100–104^.

### Cloning methods

The oligonucleotide primers (TableX)were designed based on annotated genes using goldenbraid online primer design tool for domestication and cloning into pUPD2 vector from goldenbraid 2.0 kit^105^ (Addgene kit no. 11000000076). The template cDNA for PCR was generated from immature leaf RNA using SuperScript IV RT-PCR kit (Thermo Fisher Scientific). The full-length genes were amplified using Platinum™ SuperFi II Green PCR Master Mix (Thermo Fisher Scientific) with forward and reverse primers (**Table S7**). Where multi fragment amplification was needed for domestication, the first full length amplicon was used as a template. The amplicons were then cloned into pUPD2 vector as per goldenbraid 2.0 protocol and transformed into Stellar *Escherichia coli* (Takara Bio Inc.). The positive clones were isolated using the QIAprep Spin Miniprep Kit (Qiagen) and verified for sequence integrity by Sanger sequencing (Source Bioscience, UK). All candidate genes were assembled into a pDGB3α1 vector for plant expression with 2x35S promoter and T35S terminator. The constructs were then transformed into *Agrobacterium tumefaciens* GV3101 cells using the heat-shock method. For yeast expression, genes were assembled into a modified pAV123 (addgene #63192) with GAL1 promoter and CYC1 terminator. The cytochrome P450 reductase (CPR) from *D*. *macropodum* (*DmCPR2*) was amplified from immature leaf cDNA and cloned into pESC-Leu (addgene #20120) vector under GAL1 promoter. The *S.cerevisiae* CPR (*ScCPR* -YHR042W) was amplified from genomic DNA from CEN.PK2-1C yeast wild type strain (purchased from EUROSCARF, accession no-30000B) and cloned into pESC-Leu-pGAL1_DmCPR2 vector under GAL10 promoter. Both CPRs were cloned using In-Fusion cloning kit (Takara Bio Inc.).

### Transient expression in *N. benthamiana* and leaf disc assays

The *Agrobacterium* strains harboring gene constructs of interest were grown overnight at 28°C, 220 rpm. Cells were harvested by centrifugation at 4000 ***g*** for 10 min, and the supernatant was discarded. The pellet was resuspended in 10 ml of infiltration buffer (10 mM MES, pH 5.6, 10 mM magnesium chloride, 200 μM acetosyringone) and incubated at room temperature with gentle rocking for 1-2 h. For pooled screening, strains were pooled in equal cell density with an OD_600_ of 0.3 for each strain and infiltrated into the abaxial side of a 5 week old *N. benthamiana* leaf using a 1 ml needleless syringe. Each infiltration consisted of three biological replicates, where individual leaves of three independent plants were infiltrated. Leaves were harvested 4 days post-*Agrobacterium* infiltration for plant disc assays.

For leaf disc assays, a 10 mm diameter leaf discs from *Agrobacterium* infiltrated leaves were cut using a borer. For each biological replicate, two leaf discs were used. Each plant disc was then placed in an individual well of a 24-well plate (Corning™, cat no-3526) with 500 μl of 50 mM HEPES buffer at pH 7.5 containing 25μM noscapine as internal standard with either total alkaloid extract (1:500 dilution) or pure substrate (final concentration of 25 μM) was added. Plates were closed with lid and sealed in parafilm to avoid evaporation and incubated overnight (16–18 h) in the growth room. After incubation, discs were collected, placed in a 2 ml safe-lock tube with 3 mm tungsten beads, and snap-frozen in liquid nitrogen. The discs were homogenized using a Tissue-Lyser II (Qiagen) at 25 Hz for 30-40 seconds. Metabolites were extracted from the powdered tissue with 500 μl of 80% methanol in water on a tabletop vortexer for 15 min at room temperature. The samples were then centrifuged at 16 000 rpm for 10 min to pellet cell debris. 500 μl of extract was transferred to a new 1.5 mL tube and repeated centrifugation. 100 μl the extract was transferred into analytical vials for LC-MS analysis.

### Heterologous expression in *S. cerevisiae*

The yeast strain (yss05) overexpressing truncated *HMG1* (*tHMG1*) was used as the background strain ^106^. The *DmCPR2* and *ScCPR* gene cassette was amplified as a single fragment from pESC-Leu construct and integrated near the *ARS308* locus using CRISPR/cas9 approach as described before^106^ to generate strain yss06. To test P450 activities, Dm_H1_g15271.t1 and Dm_H1_g15275.t1 constructs were transferred to yss06 background, to generate yss22 and yss23 respectively, using chemical transformation method^107^. The positive colonies were selected on Synthetic Complete (SC) selective medium lacking uracil (SC-URA) with glucose (2% w/v) as carbon source. Three independent positive colonies were inoculated on 2 mL SC-URA liquid media containing 2% galactose and 0.2% glucose in a 24-deep well plate (Fischer scientific, 12557837) and grown for 24 hrs. at 30 °C, 650 rpm. After 24 hrs media was replaced with fresh media containing 100mM HEPES pH 7.5 adjusting the starting OD1 with or without added daphhimalenine B at 6.5 µM. The cultures were grown for 24 hrs. at 30 °C, 650 rpm. The supernatant was collected and filtered through 0.2 µm PTFE filtered and transferred to LC-MS glass vials for subsequent analysis.

### LCMS analysis

LC-MS/MS runs were performed using a Vanquish Ultra-HPLC instrument interfaced to a Orbitrap Exploris™ 480 Mass Spectrometer (Thermo Fisher Scientific, Waltham, MA, USA) using an atmospheric pressure chemical ionisation (APCI) ion source in positive ion mode. 2 - μl injections were made onto an Acquity BEH C18 column (100 × 2.1 mm, 1.7 μm particle size), held at 60°C. The column was run at 0.6 ml min^−1^ with mobile phase *A* = 10 mM ammonium bicarbonate (pH 10.2) and *B* = MeOH. The gradient used was as follows: initial isocratic 2% B; 0.2–0.5 min. linear to 40% B; 0.5–5.3 min. linear to 95% B; 5.3–5.8 min. isocratic 95% B; 5.8–5.9 min. return to initial conditions and hold until 6.4 min. For untargeted analysis, LC-MS data was processed using XCMS online tool (https://xcmsonline.scripps.edu)^50^ and chromatograms were generated using R (v4.3.1) by exporting peak data through Xcalibur software (v4.1) in .csv format.

### Reaction scale-up and product purifications

The reaction scale up was done using leaf disc assay. For transient expression with target gene, 5 weeks old *N. benthamiana* plants (n=10) were used for *Agrobacterium* infiltration. The leaf discs were cut out four days after infiltration as described before. For the reaction scale up, approximately 4.5 mg of partially pure daphhimalenine B was dissolved in 120 mL of 50 mM HEPES buffer at pH 7.5 to make the reaction buffer stock. From the stock, 30 mL each was distributed to four square petri dishes. About 55-60 leaf discs were added to each plate and spread evenly in a single layer to immerse in the buffer. Plates were then closed with lid and sealed in parafilm to avoid evaporation and incubated for 48 hrs in the growth room. The buffer was collected and extracted thrice with equal volume (120 mL) of chloroform in a separating funnel to collect the chloroform fraction. The chloroform was evaporated to dryness using a rotary evaporator. The sample was resuspended in 80% methanol. The purification was carried out using Interchim Puriflash 4250 prep HPLC system linked to an Advion Expression S Compact Mass Spec (CMS). The extract was purified using a semi-preparative column (Waters XBridge C18 5µm column of size 10 x 150 mm) and eluted with a gradient of ammonium bicarbonate buffer pH = 10.2 and methanol at 4.5 mL/min flow rate. The gradient used as follows: 0-3 min 2% B, 3-12 min linear to 50% B, 12-24 min isocratic 50% B, 24-35 min linear to 60% B, 35-40 min hold at 60% B,40-55 min linear to 95% B, 55-58 min isocratic 95% B, 58-60min returning to initial conditions and hold until 65 min. The fractions were collected in 20 mL glass vials and run on Orbitrap Exploris™ 480 Mass Spectrometer as described above to confirm purity. The structures were elucidated using ^1^H, ^13^C 1D and 2D NMR methods.

### Structural elucidation by NMR

^1^H and ^13^C NMR spectra were recorded on a 700 MHz Bruker Avance Neo700 spectrometer. Chemical shifts (ppm) were recorded with tetramethylsilane or residual solvent peaks as the internal reference standard. Multiplicities are given as: s (singlet), br s (broad singlet), d (doublet), t (triplet), dd (doublet of doublets), dt (double of triples), ddd (doublet of doublet of doublets), ddt (doublet of doublet of triplets) or m (multiplet). The number of protons (*n*) for a given resonance is indicated by *n*H and coupling constants are reported as a *J* value in Hz.

### Statistical Information

Paired transcriptome and metabolome data from *D. macropodum* plant F^16^ was used for metabolite to gene correlation analysis. The normalised expression levels of genes [log2(TPM+1)] and content level of yuzurimine (as blank corrected percentage) were used to perform Spearman’s correlation coefficients (*r*) using R studio (R version 4.3.1). The genes were filtered for pfam protein families annotated as cytochrome P450, Transferase family, 2OG-Fe(II) oxygenase superfamily and dioxygenase family. Pearson’s correlation coefficients were calculated using log2(TPM+1) against the bait gene *Dm_g8422.t1*. For statistical comparison of peak areas to determine substrate consumption and product formation, we used two tailed Student’s *t*-tests with *n* = 3 to compare reactions against empty vector controls. Error bars show standard deviation.

## Data Availability

The *Daphniphyllum macropodum* genome assembly will be available from NCBI Genomes (haplotype 1: BioProject PRJNA1462275, Accession JBXYWJ000000000; haplotype 2: BioProject PRJNA1462274, Accession JBXYWK000000000). The genome assembly and annotations will be available on FigShare (10.6084/m9.figshare.32608323). The genomic and transcriptomic raw reads will be available from the NCBI sequence read archive (BioProject PRJNA1462175, accessions SRR38384357-SRR38384390). All data will be released upon publication of the article.

## Acknowledgements

The plasmid pHREAC was a gift from George Lomonossoff (Addgene plasmid #134908). We are grateful for computational support from the University of York High-Performance Computing service, Viking, and the Research Computing team. We acknowledge the University of York Biology Technology Facility, the Centre of Excellence in Mass Spectrometry, the Biology Horticultural Facility and the Chemistry NMR facility in conducting this research. We are grateful for advice from Mohamed Kamileen about the leaf disc assays. We thank the Yorkshire Arboretum and Castle Howard for access to the plant material. We thank David Nelson for providing the names of the cytochrome P450s. The work was supported through a UKRI Future Leaders Fellowship awarded to BRL (MR/S01862X/1 and MR/X010260/1). BAR was supported by a BBSRC White Rose DTP studentship (BB/T007222/1).

## Author contributions

S.S. performed correlation analysis, selected gene candidates, cloned genes, conducted and analysed enzyme assays (*N. benthamiana* leaf discs and yeast), performed scale-up reactions and compound isolations, and wrote the manuscript. B.A.R. and S.L. isolated and purified alkaloids from *D. macropodum*. B.A.R., L.W. and W.P.U. characterised alkaloids. D.J.T. grew *D. macropodum* and isolated nucleic acids. S.J.S. annotated genome assembly and performed phylogenetic analysis. T.R.L. supported metabolite analysis. B.R.L. managed the project, performed data analysis and wrote the manuscript.

## Ethics declaration

The authors declare no competing interests.

## Supplementary Information

Supplementary Figures S1-S24 Supplementary Tables S1-S7 Supplementary References

