## Supplementary Information for "*Daphniphyllum macropodum* genome assembly and multiplexed screening unveils alkaloid modifying enzymes"

For

### Supplementary Figures

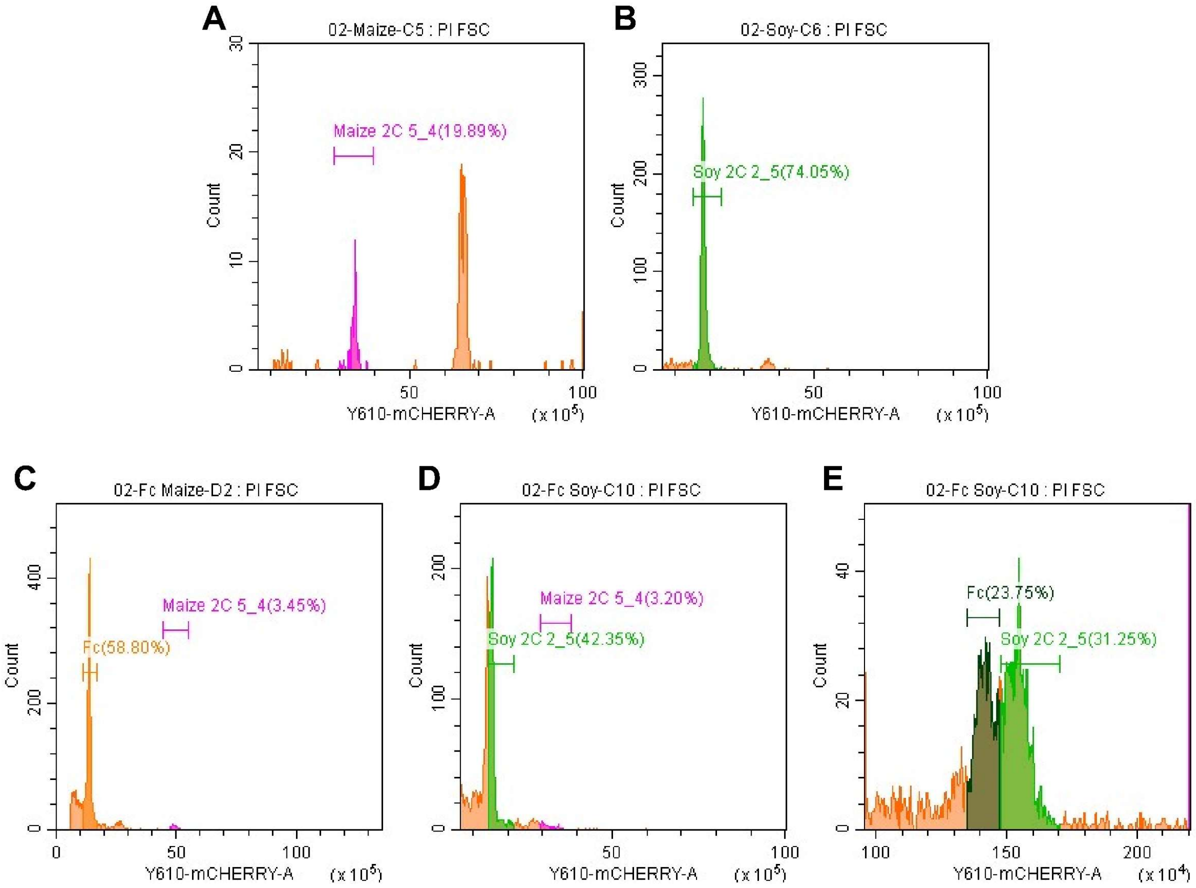

**Supplementary Fig. S1**. **Flow cytometry to determine *Daphniphyllum macropodum* genome size. A.** Maize (*Zea mays* CE-777) sample, showing 2C peak (5310 Mbp). Larger peak represents somatic whole genome duplication. **B.** Soy (*Glycine max* polanka) sample, showing C2 peak (2445 Mbp). **C.** *D. macropodum* sample (plant Fc, orange) compared to maize sample (pink). **D.** *D. macropodum* sample (plant Fc, orange) compared to soy sample (green). **E.** Zoomed in comparison of *D. macropodum* sample (plant Fc, brown) compared to soy sample (green). The X-axis represents DNA content via propidium iodide (PI), measured through the Y610-mCHERRY-A channel.

**A**

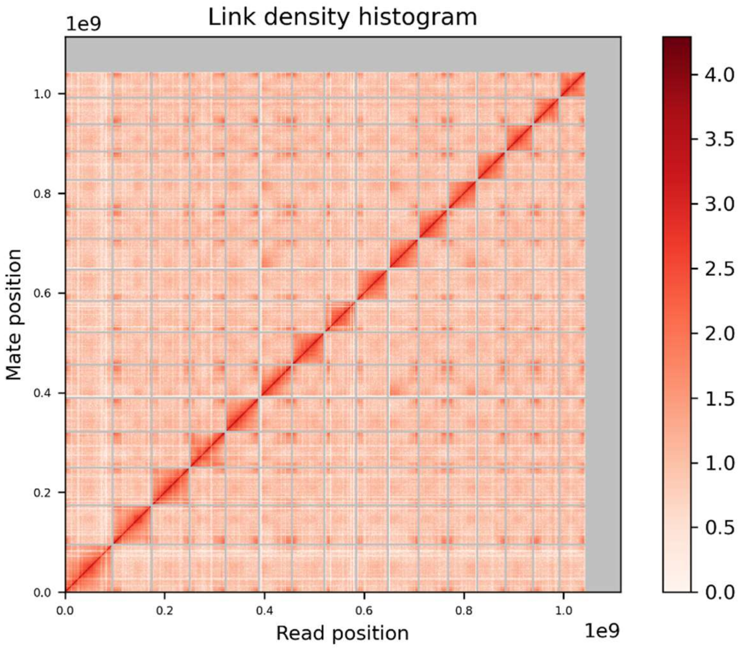

**B**

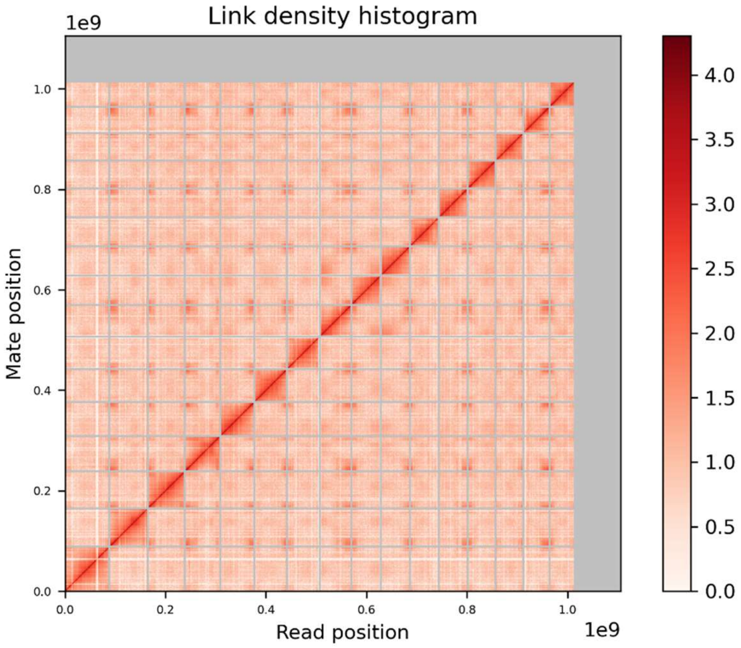

**Supplementary Figure S2. HiRise Scaffolding. A.** Link density histogram for haplotype 1. **B.** Link density histogram for haplotype 2.

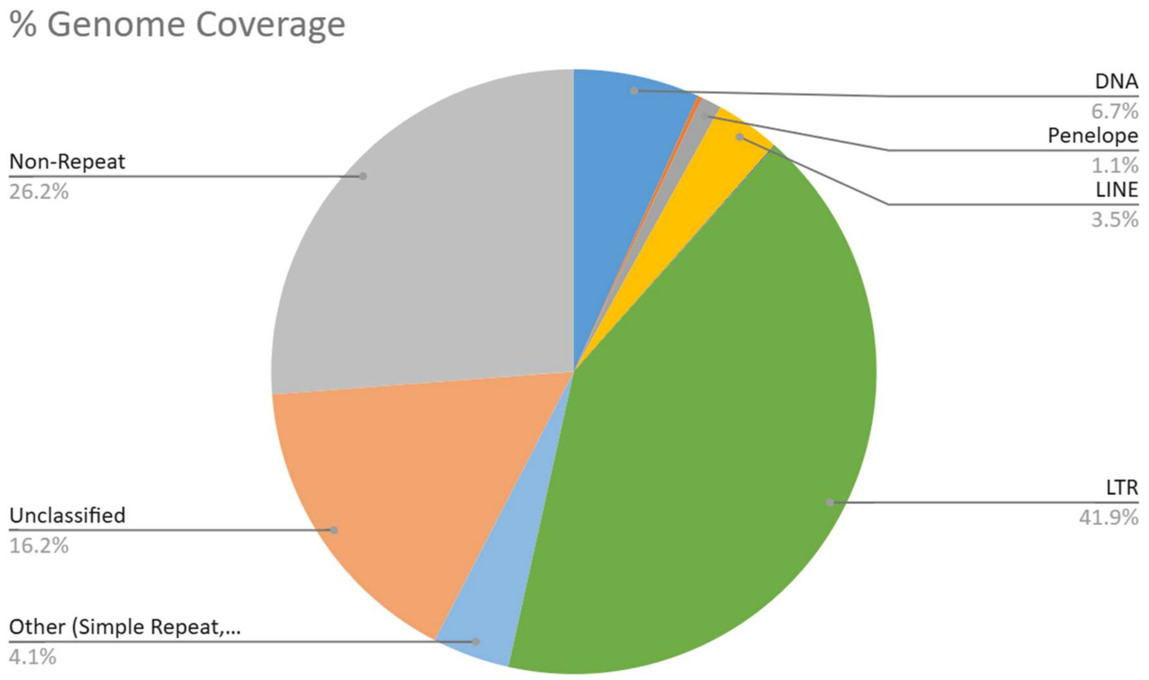

**Supplementary Figure S3.** Density information of repeats/transposable elements (haplotype 1). See **Table S2** for further details.

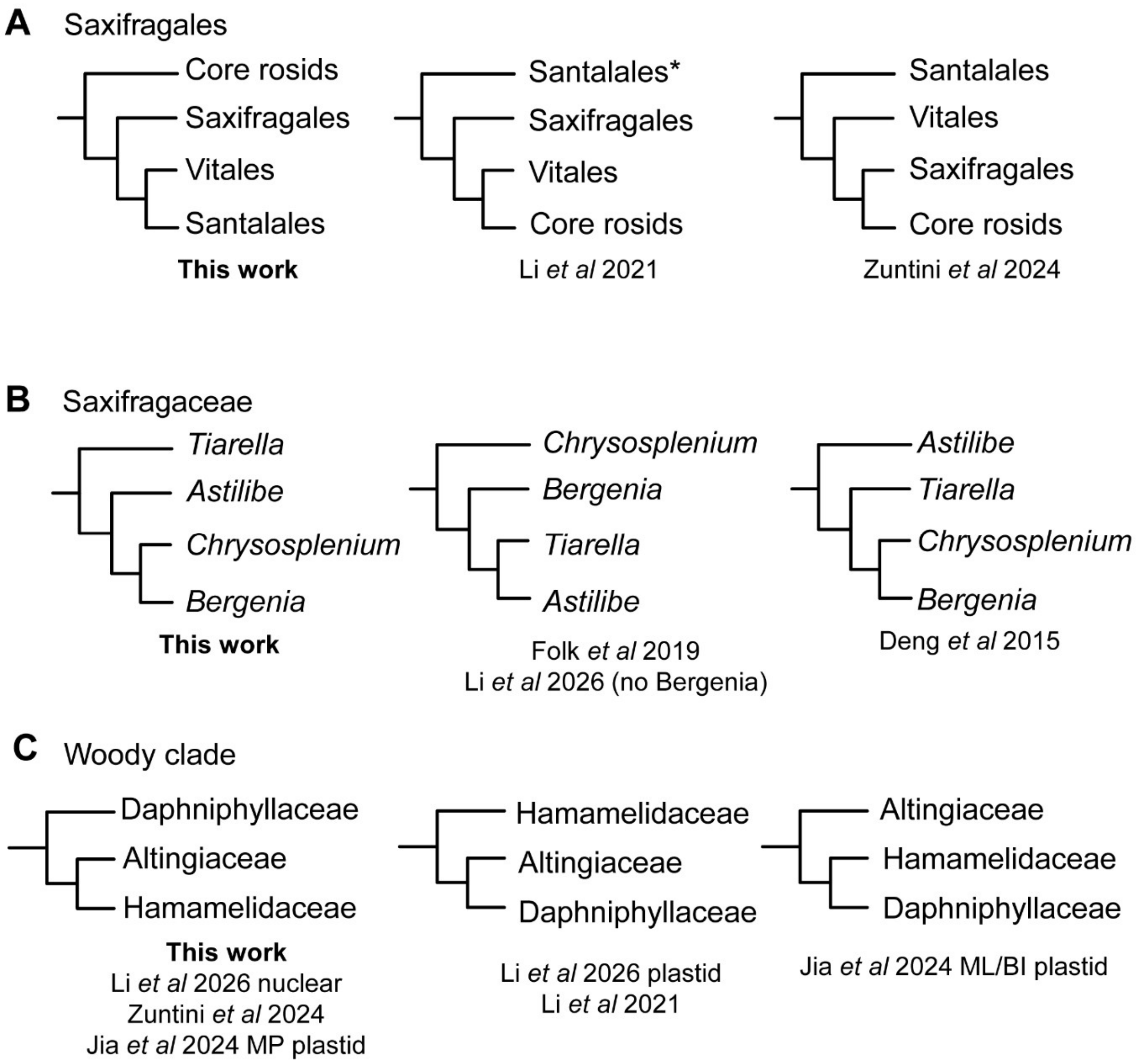

**Supplementary Figure S4. Phylogenetic relationships in the Saxifragales. A.** Position of Saxifragales relative to superrosids and related clades, rooted on core asterids. * = in Li *et al* 2021 Santalales was recovered as sister to core asterids but depicted here as sister to superrosids due to adjusted rooting. **B.** Inferred relationships between selected genera in Saxifragaceae. For Li *et al* 2026 no Bergenia sample was included in the inference. **C.** Topology of three families in the Saxifragales woody clade. References: Li *et al* 2021[1], Zuntini *et al* 2024[2], Folk *et al* 2019[3], Li *et al* 2026[4], Deng *et al* 2015[5], Jia *et al* 2024[6].

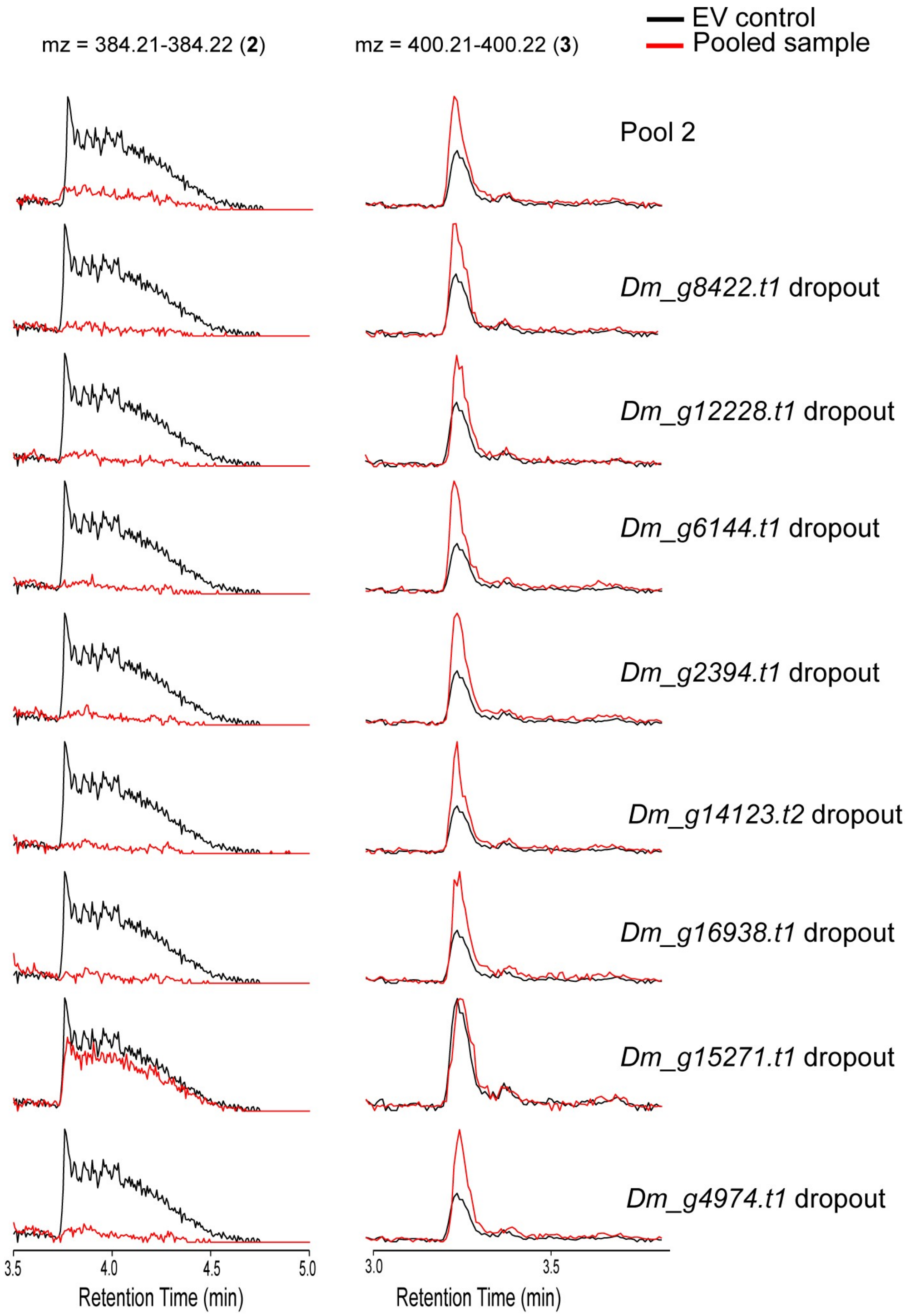

**Supplementary Figure S5**. HR-LC-MS/MS extracted ion chromatograms of gene dropout experiment to identify the active enzyme. The extracted ion chromatograms for *Dm_H1_g15271.t1* drop out shows no drastic change compared to control suggesting its involvement in the activity. See Fig. 5F and 5G for corresponding bar charts.

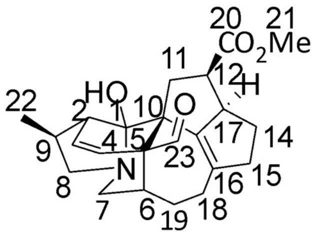

daphhimalenine B

Chemical Formula: $C_{23}H_{29}NO_4$

Exact Mass: 383.21

| C no. | | Daphhimalenine B (literature) $\delta_C$ (1% TFA) | Daphhimalenine B (experimental) $\delta_C$ |
|---|---|---|---|
| **1** | C | 103.3 | 99.5 |
| **2** | CH | 46.5 | 47.7 |
| **3** | CH/CH/$CH_2$ | 127.0 | 127.9 |
| **4** | CH/CH/C | 130.1 | 130.0 |
| **5** | C | 48.3 | 48.9 |
| **6** | CH | 38.2 | 39.3 |
| **7** | $CH_2$ | 54.4 | 53.7 |
| **8** | $CH_2$ | 61.6 | 62.6 |
| **9** | CH | 32.4 | 34.2 |
| **10** | C | 56.4 | 57.2 |
| **11** | $CH_2$ | 41.7 | 41.1 |
| **12** | CH | 43.0 | 43.1 |
| **13** | CH | 58.2 | 58.1 |
| **14** | $CH_2$ | 29.8 | 29.8 |
| **15** | $CH_2$ | 43.2 | 43.4 |
| **16** | C | 140.4 | 137.9 |
| **17** | C | 142.0 | 144.1 |
| **18** | $CH_2$ | 25.8 | 26.1 |
| **19** | $CH_2$ | 25.0 | 26.8 |
| **20** | C | 175.8 | 176.0 |
| **21** | $CH_3$ | 51.5 | 51.5 |
| **22** | $CH_3$ | 18.2 | 18.0 |
| **23** | CH | 206.1 | 209.0 |

**Supplementary Fig. S6**. **$^{13}$C NMR data for daphhimalenine B (2).** Comparison between literature values $^{13}$C (125 MHz) and experimental $^{13}$C (175 MHz) NMR data of daphhimalenine B in $CDCl_3$ [7].

| C | | daphhimalenine B (experimental) $\delta_H$ J (Hz) | daphhimalenine B (literature) in 1% TFA $\delta_H$ J (Hz) |
|---|---|---|---|
| **1** | C | | |
| **2** | CH | 2.82 – 2.65 m | 3.05 m |
| **3** | **$CH_2$/CH** | 5.74 dd 10.5, 3.0 | 5.85 dd 10.5, 3.0 |
| **4** | **CH/CH** | 6.03 dd 10.5, 2.0 | 6.16 dd 10.5, 2.0 |
| **5** | C | | |
| **6** | CH | 2.62 – 2.50 m | 2.18 m |
| **7** | $CH_2$ | 3.16 – 3.05 m<br>2.93 dd 12.0, 7.0 | 3.64 m<br>3.28 m |
| **8** | $CH_2$ | 3.59 – 3.44 m<br>2.30 dd 11.0, 5.5 | 4.19 dt 12.0, 3.5<br>2.69 dd, 12.0, 3.5 |
| **9** | CH | 2.82 – 2.65 m | 3.18 m |
| **10** | C | | |
| **11** | $CH_2$ | 2.53 dd 15.0, 9.5<br>2.25 dd 15.0, 4.0 | 2.55 dd 15.5, 9.5<br>2.33 dd 15.5, 3.0 |
| **12** | CH | 3.16 – 3.05 m | 3.08 m |
| **13** | CH | 3.75 – 3.68 m | 3.87 br s |
| **14** | $CH_2$ | 1.99 – 1.92 m<br>1.43 – 1.34 m | 1.97 m<br>1.27 m (overlapped) |
| **15** | $CH_2$ | 2.82 – 2.65 m<br>2.42 d 7.5 | 2.64 m<br>2.37 d 7.5 |
| **16** | C | | |
| **17** | C | | |
| **18** | $CH_2$ | 1.99 – 1.92<br>1.68 – 1.60 | 1.94 m<br>1.70 m |
| **19** | $CH_2$ | 2.62 – 2.56 m<br>2.17 – 2.11 m | 2.62 m<br>2.23 d 16.5 |
| **20** | C | | |
| **21** | $CH_3$ | 3.63 s | 3.61 s |
| **22** | $CH_3$ | 1.18 d 7.5 | 1.26 d 7.5 |
| **23** | $CH_2$/CH | 10.33 s | 10.37 s |

**Supplementary Fig. S7**. **$^1$H NMR data for daphhimalenine B (2).** Comparison between experimental values for $^1$H NMR data of daphhimalenine B (700 MHz) in $CDCl_3$, and literature values for daphhimalenine B in $CDCl_3$ in 1% TFA [7].

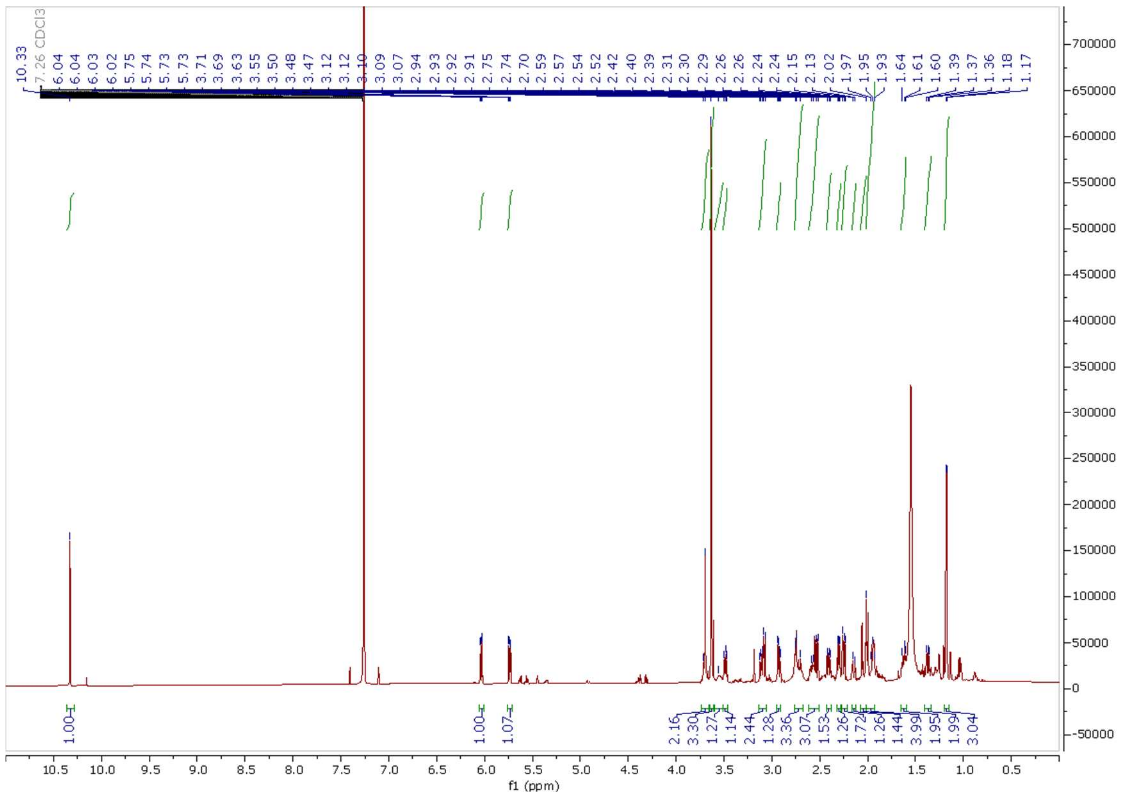

**Supplementary Figure S8. $^{1}$H NMR spectrum of daphhimalenine B (2) in $CDCl_3$.**

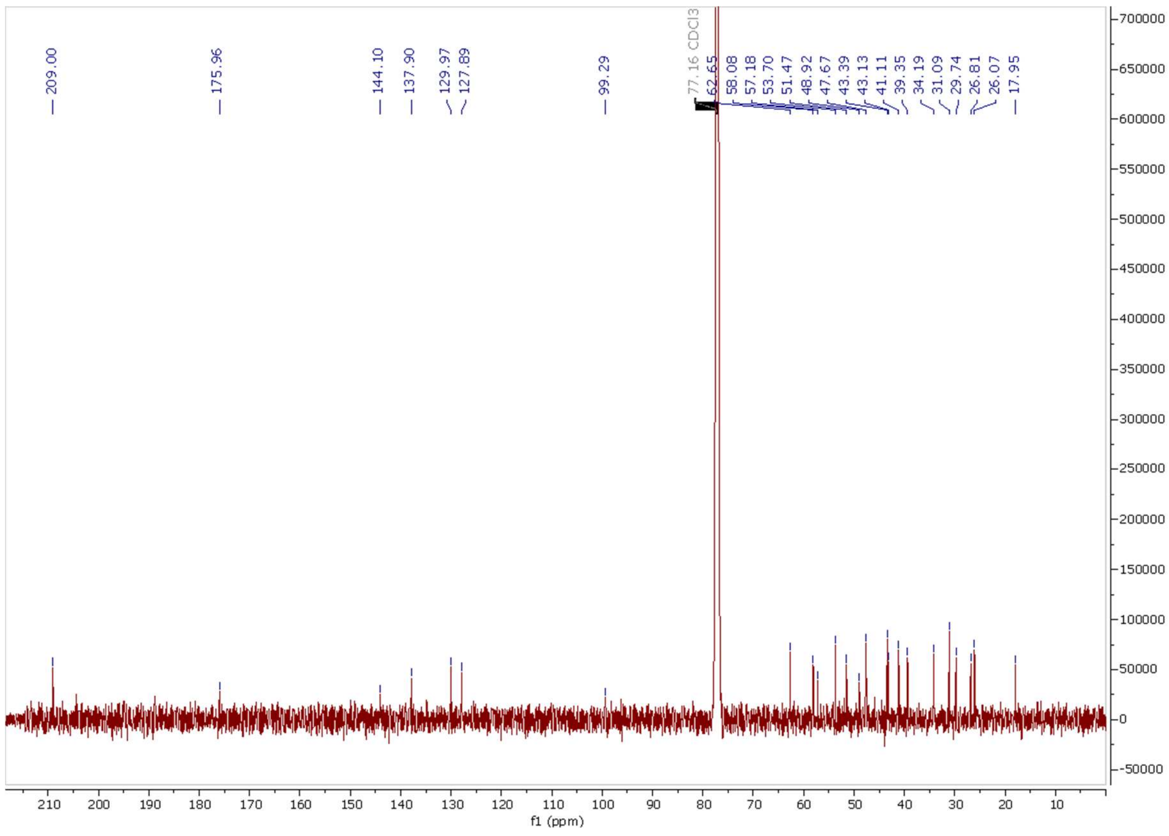

**Supplementary Figure S9. $^{13}$C NMR spectrum of daphhimalenine B (2) in $CDCl_3$.**

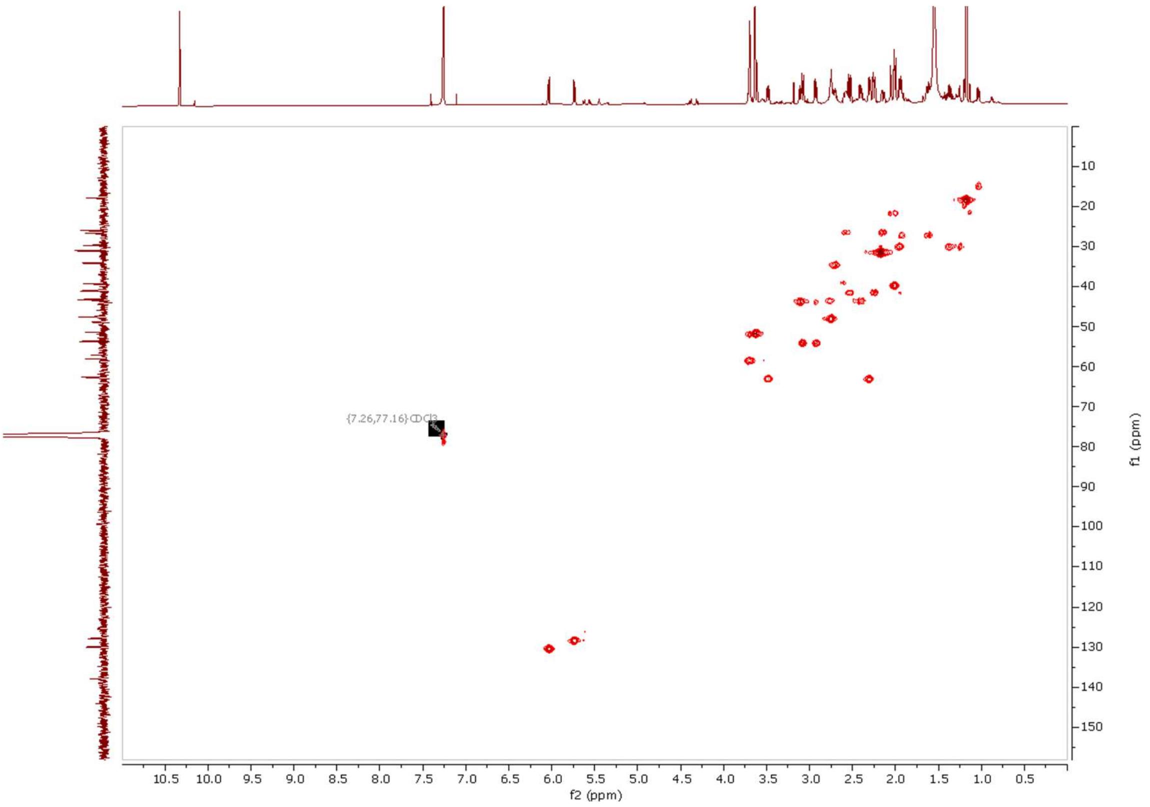

**Supplementary Figure S10. HMQC NMR spectrum of daphhimalenine B (2) in $CDCl_3$.**

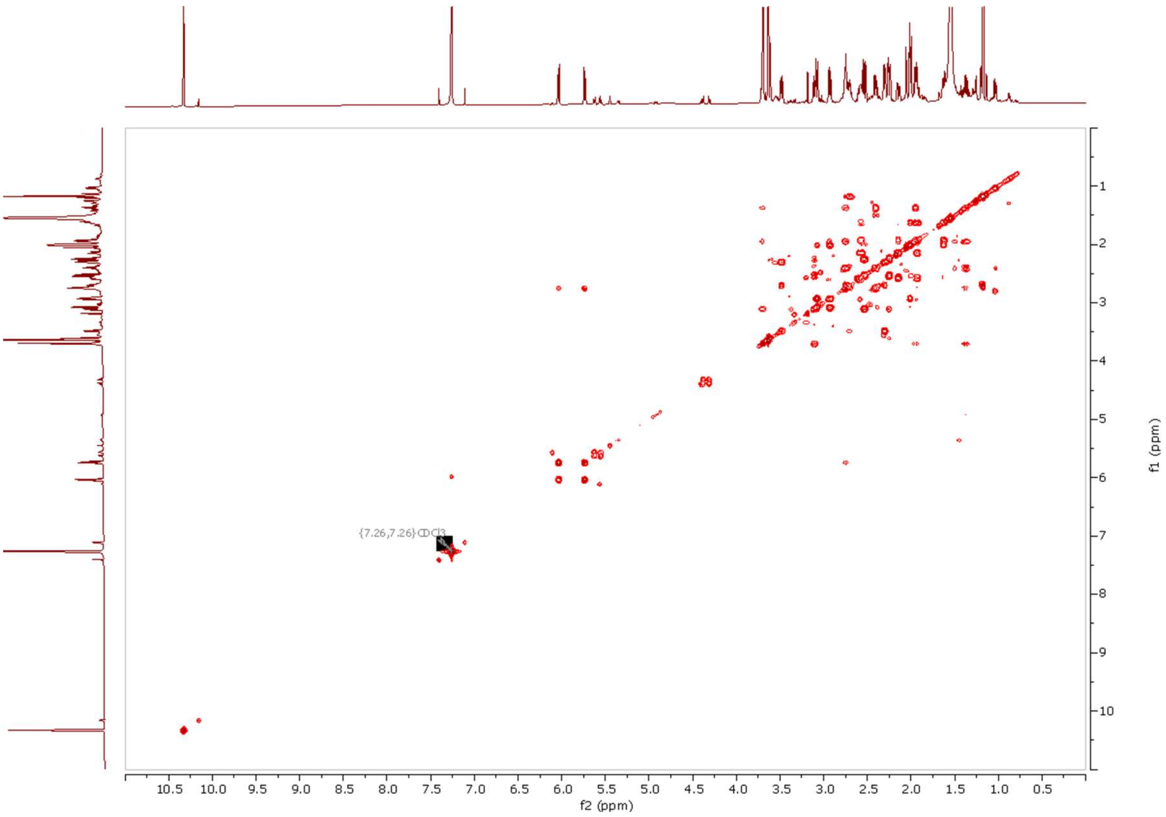

**Supplementary Fig. S11. COSY NMR spectrum of daphhimalenine B in $CDCl_3$.**

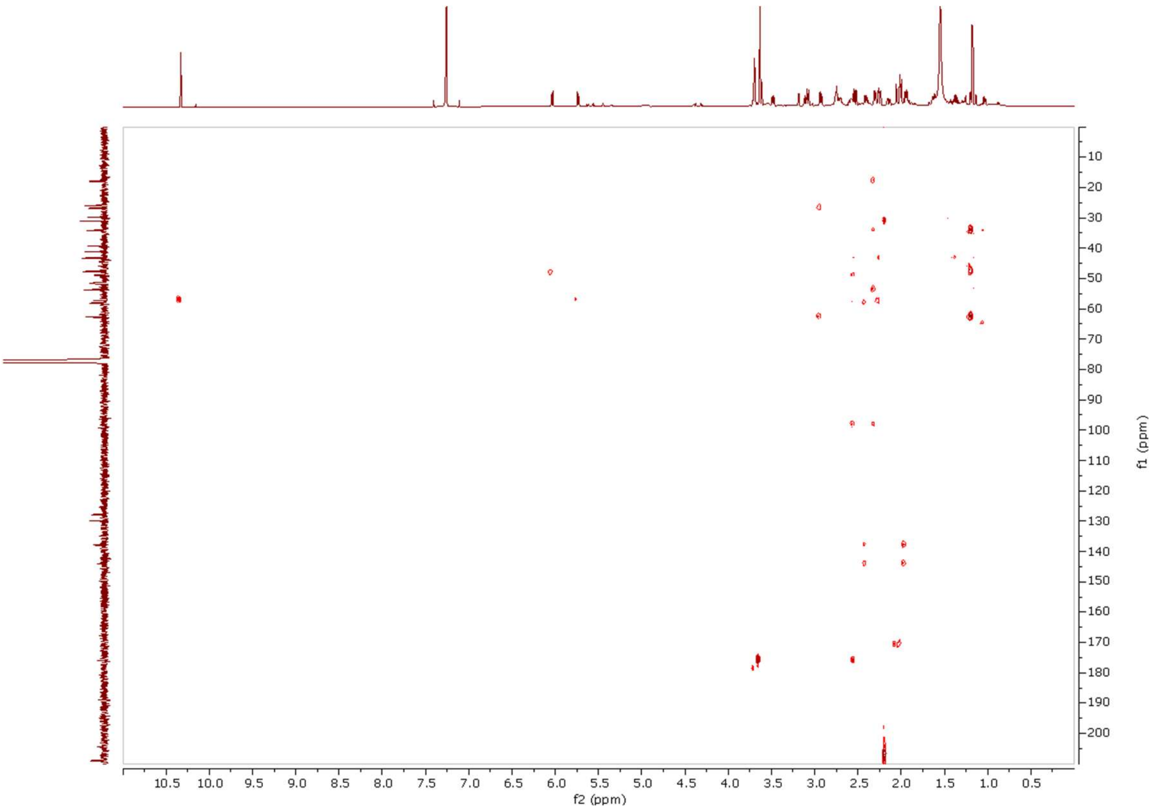

**Supplementary Fig. S12. HBMC NMR spectrum of daphhimalenine B (2) in $CDCl_3$.**

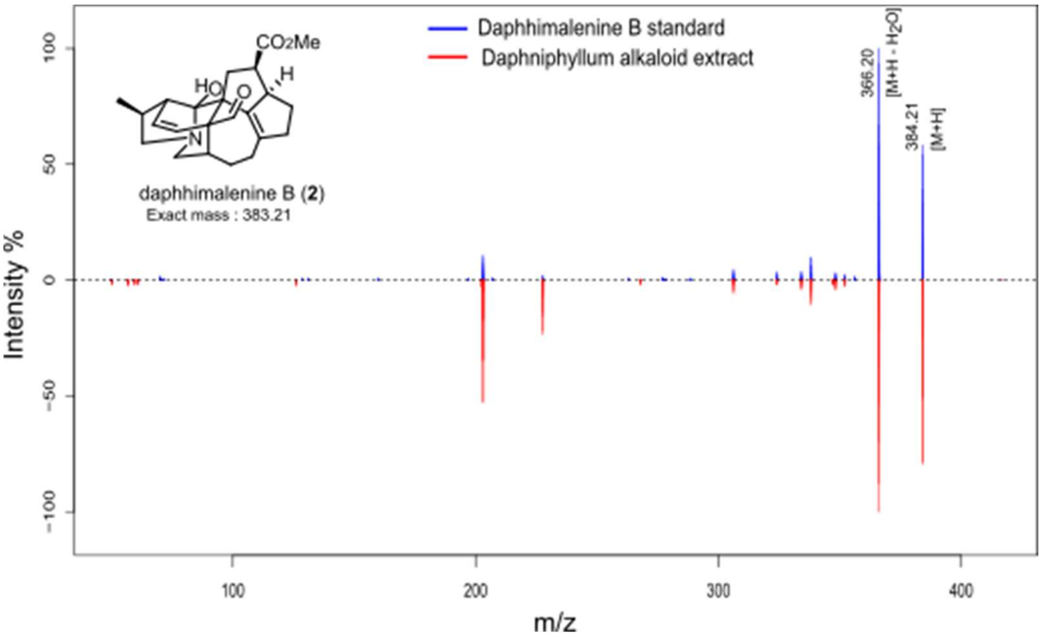

**Supplementary Figure S13. Daphhimalenine B (2) MS$^2$ data.** Standard data (top) matches the observed substrate in the alkaloid extract (bottom).

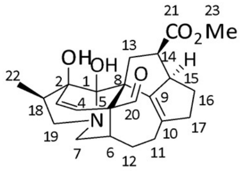

yuzurimine C
Chemical Formula: $C_{23}H_{29}NO_5$
Exact Mass: 399.2046

| C no. | | Literature $\delta_C$ | Experimental $\delta_C$ | Literature $\delta_H$ | Experimental $\delta_H$ |
|---|---|---|---|---|---|
| **1** | C | 91.6 | 92.5 | | |
| **2** | C | 80.1 | 80.2 | | |
| **3** | CH | 125.0 | 125.5 | 5.61 d, *J* = 10.0 Hz | 5.68 d, *J* = 10.5 Hz |
| **4** | CH | 132.5 | 132.4 | 6.20 d, *J* = 10.0 Hz | 6.26 d, *J* = 10.5 Hz |
| **5 (8)** | C | 48.7 | 48.5 | | |
| **6** | CH | 36.0 | 36.4 | | 2.17–2.10 m |
| **7** | $CH_2$ | 49.1 | 50.0 | | 3.45 dd, *J* = 14.0, 8.0 Hz<br>2.77 dd, *J* = 14.0, 8.0 Hz |
| **8 (5)** | C | 55.2 | 55.5 | | |
| **9** | C | 136.9 | 137.3 | | |
| **10** | C | 125.0 | 142.9 | | |
| **11** | $CH_2$ | 25.6 | 25.6 | | 2.46–2.39 m<br>2.31–2.23 m |
| **12** | $CH_2$ | 24.8 | 25.2 | | 2.08–1.89 m<br>1.68–1.41 |
| **13** | $CH_2$ | 42.4 | 36.0 | | 3.23–3.11 m<br>1.85 dd, *J* = 14.5, 7.6 Hz |
| **14** | CH | 55.9 | 43.1 | | 3.23–3.11 m |
| **15** | CH | 43.0 | 56.3 | | 3.88–3.81 m |
| **16** | $CH_2$ | 29.6 | 29.9 | | 2.08–1.89 m<br>1.38–1.29 m |
| **17** | $CH_2$ | 35.5 | 42.7 | | 2.68–2.62 m<br>2.31–2.23 m |
| **18** | CH | 40.6 | 41.4 | | 2.23–2.17 m |
| **19** | $CH_2$ | 56.8 | 57.2 | | 3.23–3.11 m<br>2.17–2.10 m |
| **20** | CH | 207.0 | 207.3 | 9.99 s | 10.09 s |
| **21** | C | 175.6 | 175.9 | | |
| **22** | $CH_3$ | 11.4 | 12.4 | 1.04 d 6.0 Hz | 1.05 d, 7.0 Hz |
| **23** | $CH_3$ | 51.2 | 51.5 | 3.58 s | 3.63 s |

**Supplementary Fig. S14**. **$^1H$ and $^{13}C$ NMR data for yuzurimine C (3)**. Comparison between literature values $^1H$ (100MHz), $^{13}C$ (15 MHz) and experimental $^1H$ (700 MHz), $^{13}C$ (100 MHz) NMR data of yuzurimine C in $CDCl_3$ [8].

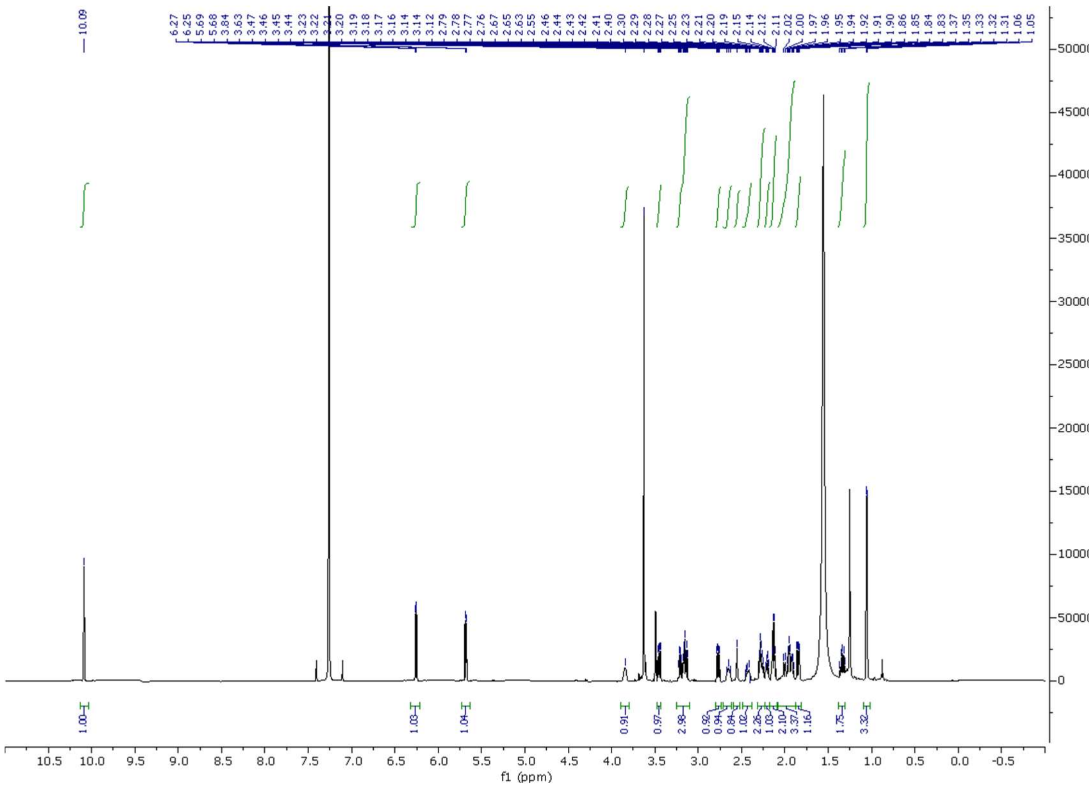

**Supplementary Figure S15. $^{1}$H NMR spectrum of yuzurimine C (3) in $CDCl_3$.**

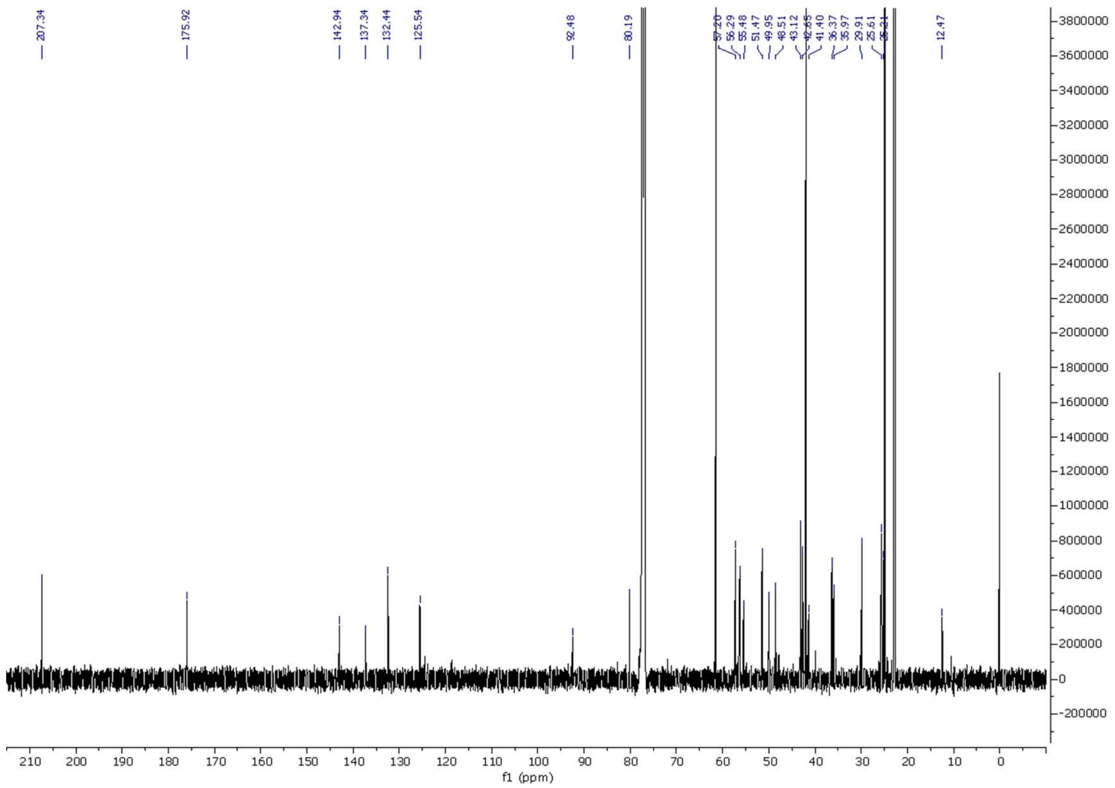

**Supplementary Fig. S16. $^{13}$C NMR spectrum of yuzurimine C in $CDCl_3$.**

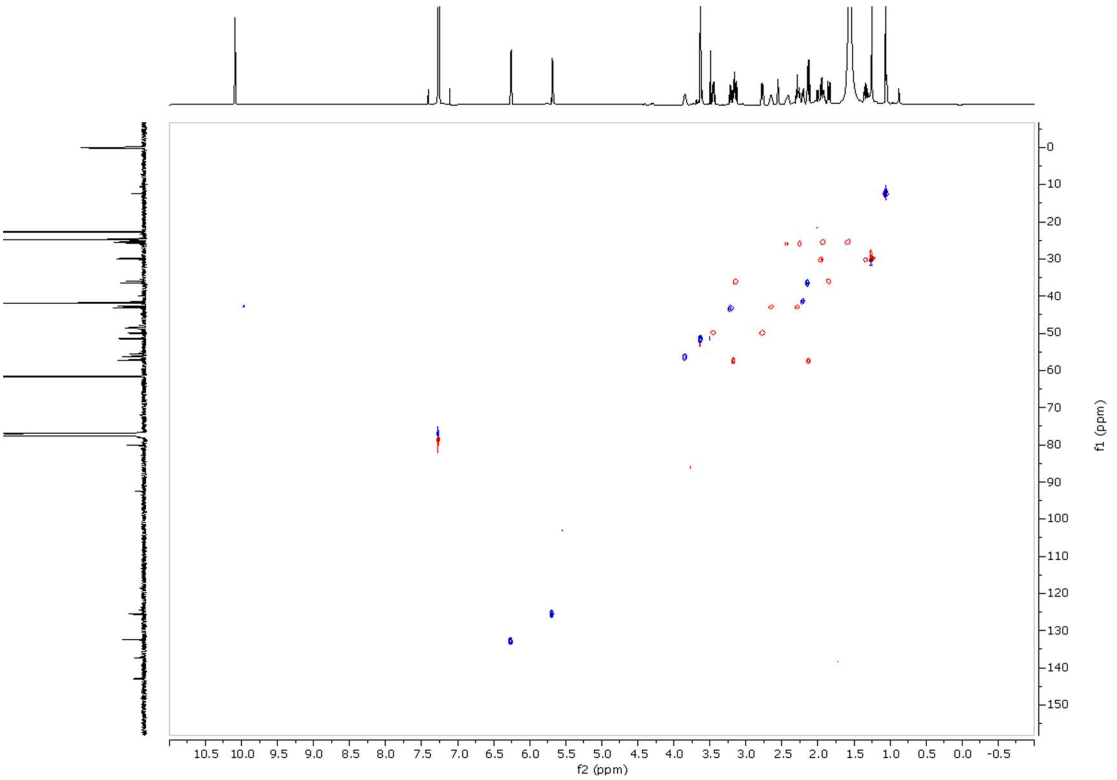

**Supplementary Fig. S17. HSQC NMR spectrum of yuzurimine C (3) in $CDCl_3$.**

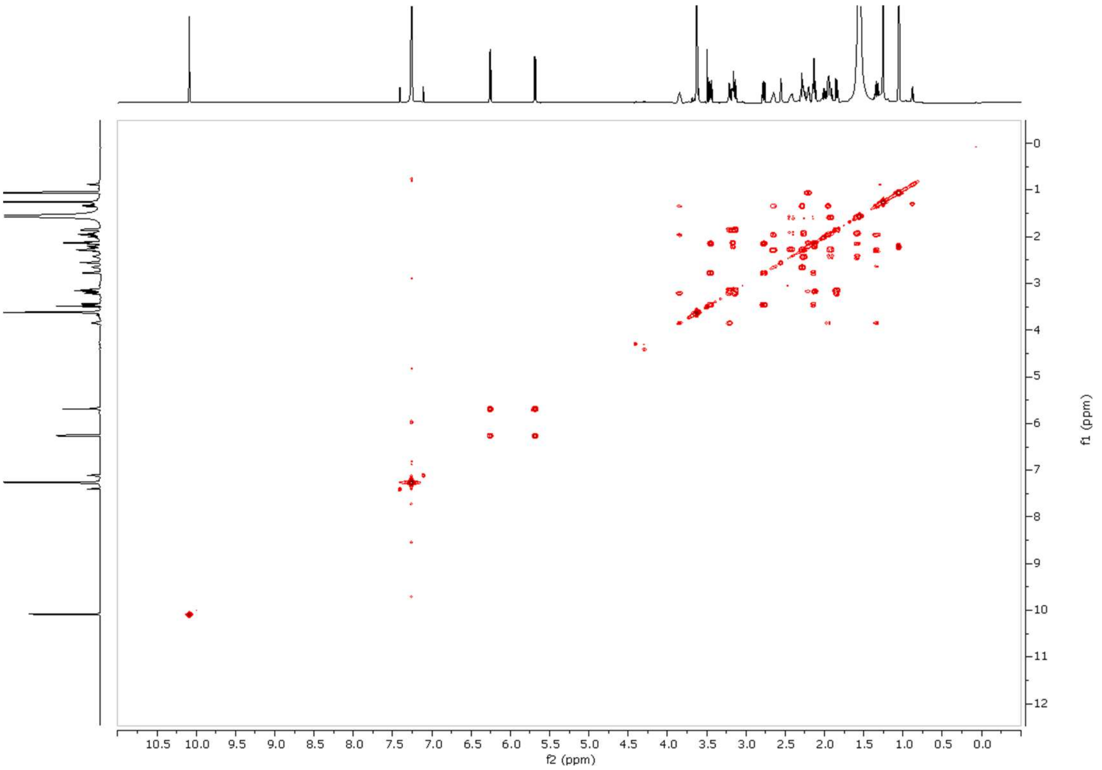

**Supplementary Fig. S18. COSY NMR spectrum of yuzurimine C (3) in $CDCl_3$.**

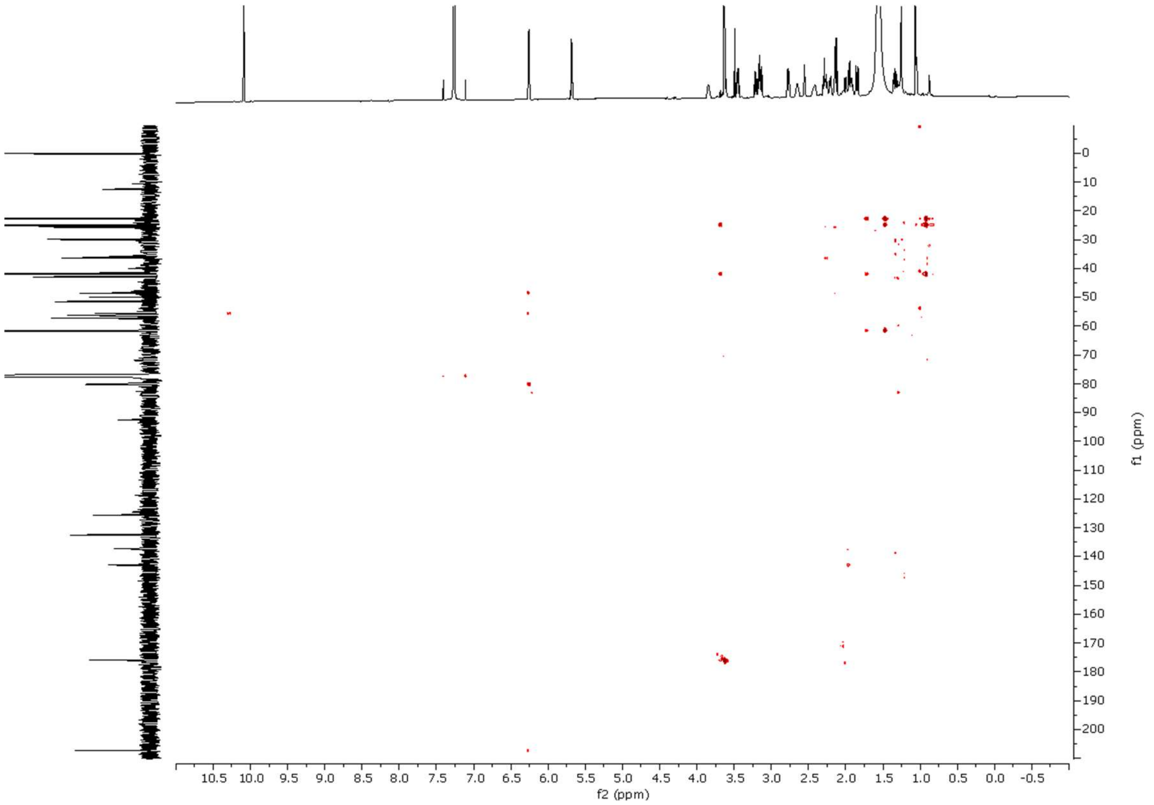

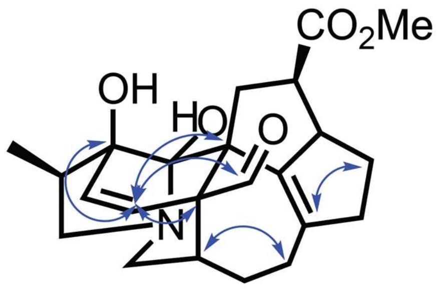

yuzurimine C (**3**)

**Supplementary Fig. S19. HMBC NMR spectrum of yuzurimine C (3) in $CDCl_3$.** Top – HMBC spectrum. Bottom – coupling connections deduced from spectrum.

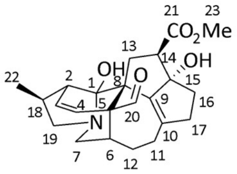

yuzurimine F
Chemical Formula: $C_{23}H_{29}NO_5$
Exact Mass: 399.2046

| C no. | | $\delta_C$ | $\delta_H$ |
|---|---|---|---|
| **1** | C | 97.1* | |
| **2** | C | 50.9 | 2.78–2.74 m |
| **3** | CH | 126.5 | 5.71 d, *J* = 10.4, 2.8 Hz |
| **4** | CH | 130.5 | 6.09 d, *J* = 10.4 Hz |
| **5 (8)** | C | 56.9 | |
| **6** | CH | 38.7 | 2.15–2.11 m |
| **7** | $CH_2$ | 50.9 | 3.30 dd, *J* = 13.4, 4.7 Hz<br>2.91 dd, *J* = 13.4, 7.6 Hz |
| **8 (5)** | C | 49.4 | |
| **9** | C | 143.9* | |
| **10** | C | 144.4* | |
| **11** | $CH_2$ | 26.0 | 2.74–2.63 m<br>2.27–2.19 m |
| **12** | $CH_2$ | 25.8 | 1.95–1.88 m<br>1.67–1.61 m |
| **13** | $CH_2$ | 38.8 | 2.51 dd, *J* = 15.0, 9.5 Hz<br>2.27–2.19 m |
| **14** | CH | 53.2 | 3.16 dd *J* = 9.5, 6.3 Hz |
| **15** | C | 96.0 | |
| **16** | $CH_2$ | 36.7 | 2.02 dd, *J* = 12.9, 5.9 Hz<br>1.84 dt, *J* = 12.9, 8.4 Hz |
| **17** | $CH_2$ | 41.3 | 3.11 dt, *J* = 15.5, 7.5 Hz<br>2.33 dd, *J* = 15.5, 8.4 Hz |
| **18** | CH | 33.1 | 2.74–2.63 m |
| **19** | $CH_2$ | 61.2 | 3.50–3.39 m<br>2.39 dd, *J* = 9.9, 7.2 Hz |
| **20** | CH | 207.4 | 10.15 s |
| **21** | C | 174.5 | |
| **22** | $CH_3$ | 16.2 | 1.17 d, *J* = 7.2 Hz |
| **23** | $CH_3$ | 51.9 | 3.69 s |

**Supplementary Fig. S20**. **$^{1}$H and $^{13}$C NMR data for yuzurimine F (4)**. $^{13}$C Peaks labelled with a star were inferred from HMBC spectrum.

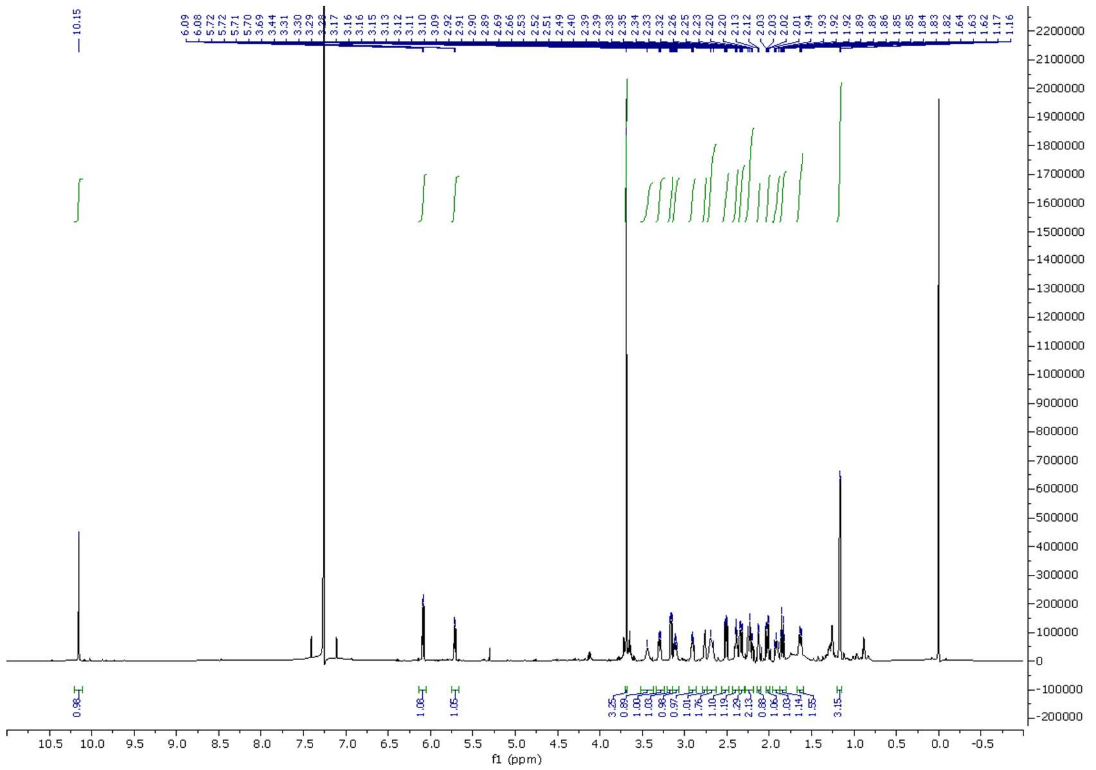

**Supplementary Fig. S21. $^{1}$H NMR spectrum of yuzurimine F (4) in $CDCl_3$.**

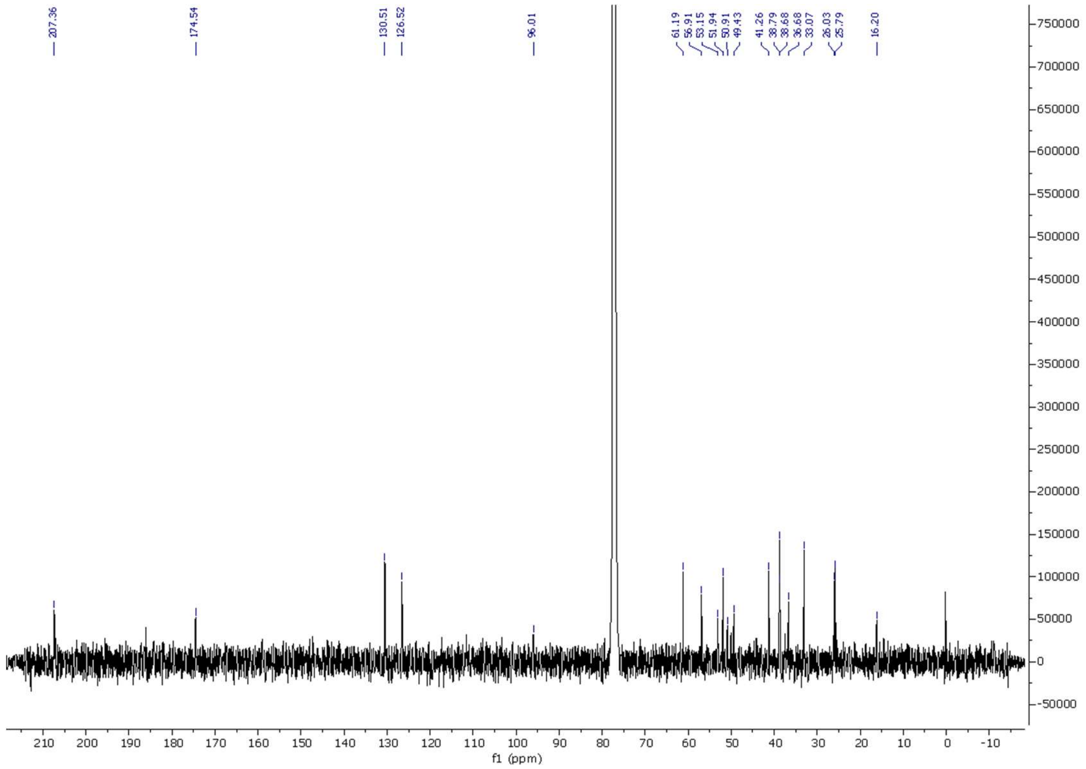

**Supplementary Fig. S22. $^{13}$C NMR spectrum of yuzurimine F (4) in $CDCl_3$.**

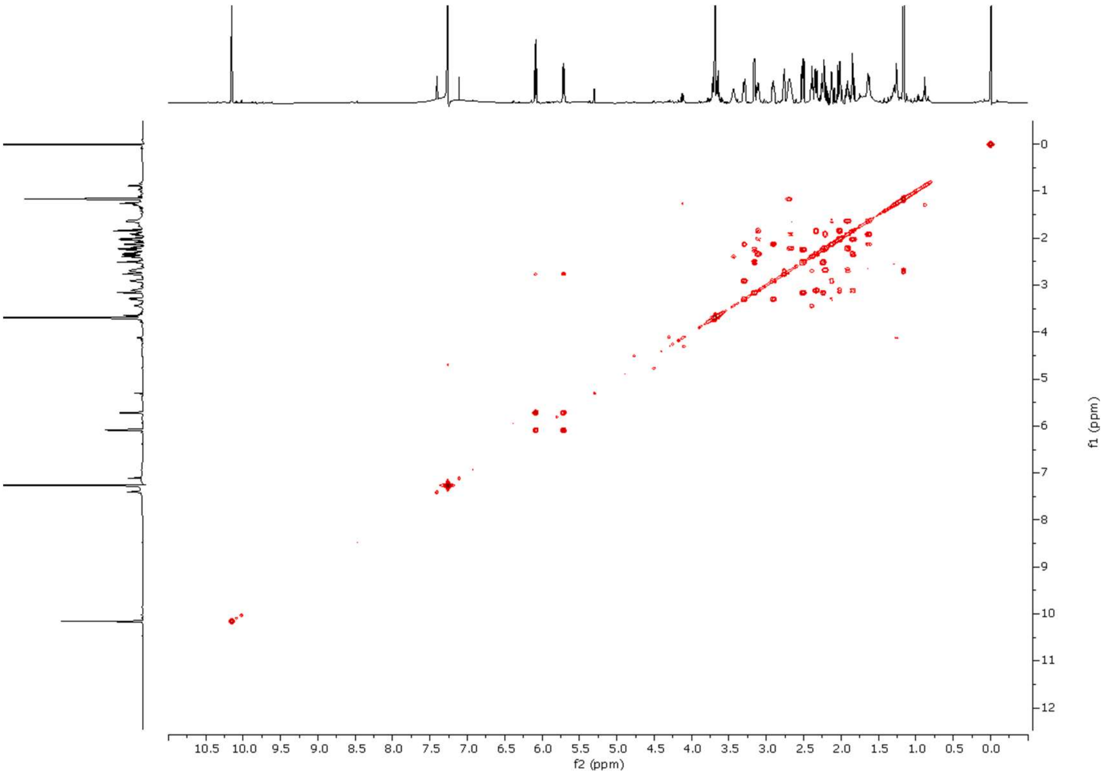

**Supplementary Fig. S23. COSY NMR spectrum of yuzurimine F (4) in $CDCl_3$.**

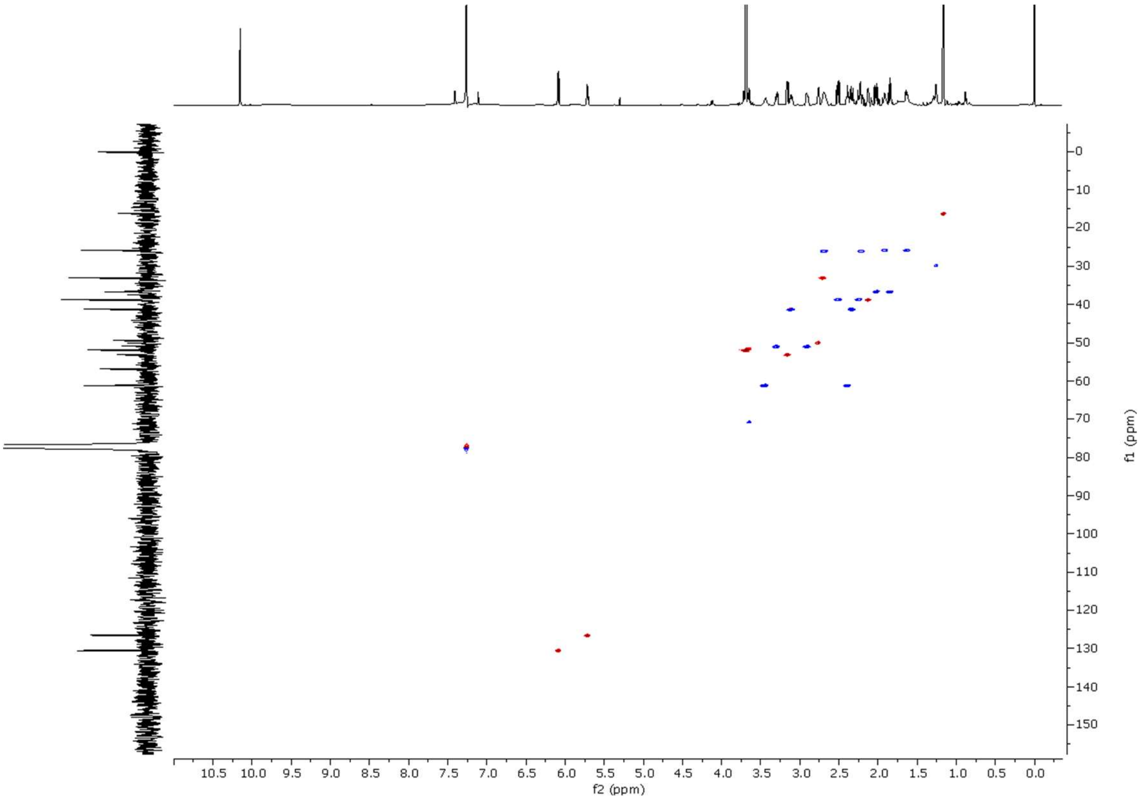

**Supplementary Fig. S24. HSQC NMR spectrum of yuzurimine F (4) in $CDCl_3$.**

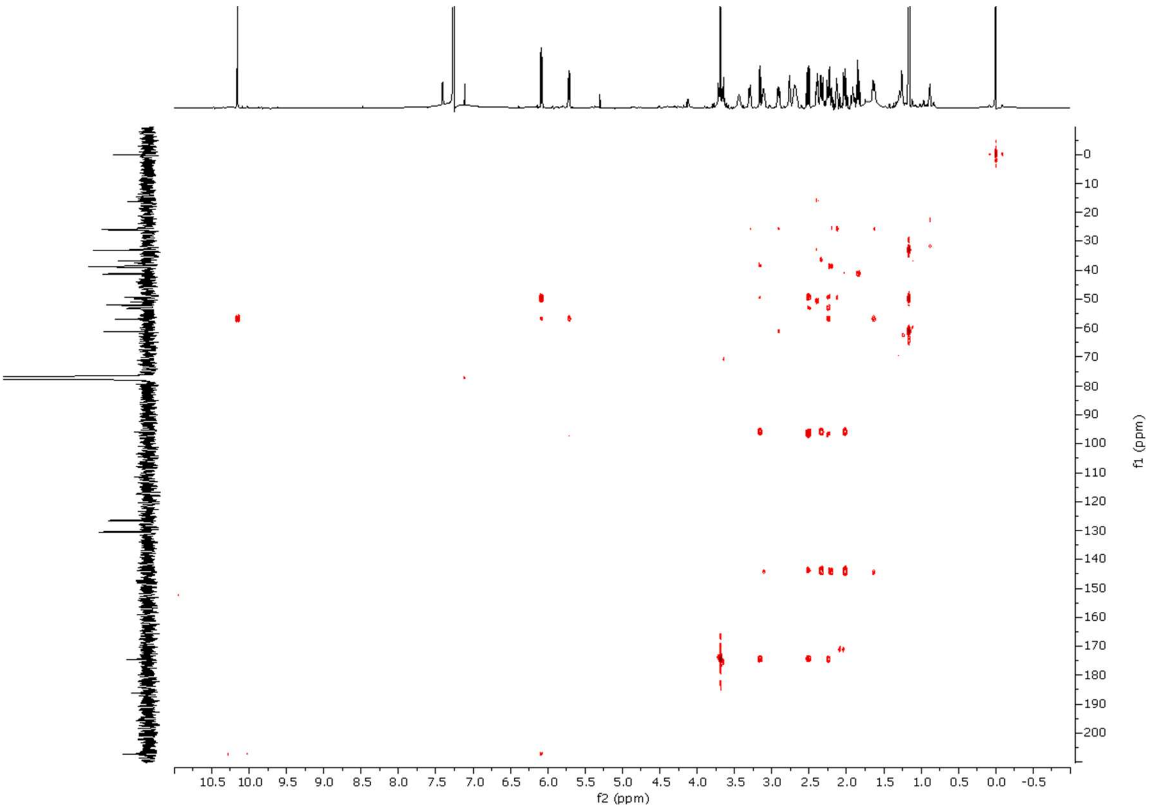

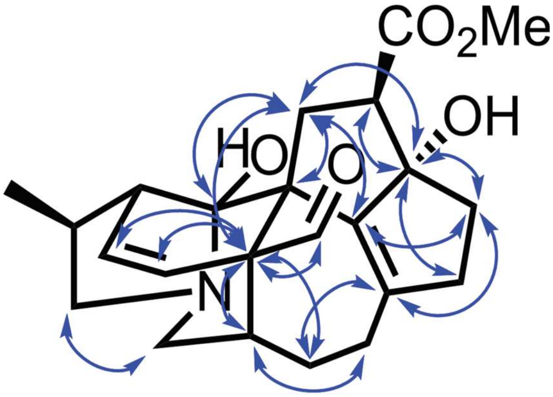

yuzurimine F (**4**)

**Supplementary Fig. S24. HMBC NMR spectrum of yuzurimine F (4) in $CDCl_3$.** Top – HMBC spectrum. Bottom – coupling connections deduced from spectrum.

### Supplementary Tables

**Table S1**. Genome assembly metrics.

| ASSEMBLY | hifiasm | HAP1 | HAP2 |
|---|---|---|---|
| Total Assembly (Mbp) | 1181.6 | 1105.76 | 1114.35 |
| Assembled to Chr. (Mbp) | - | 1014.73 (91.8%) | 1045.13 (93.8%) |
| N50 (Mbp) | 34.04 | 62.89 | 63.47 |
| L50 | 12 | 8 | 8 |
| N90 (Mbp) | 1.46 | 50.38 | 53.3 |
| L90 | 55 | 16 | 16 |
| Number of Scaffolds | 2900 | 2240 | 785 |
| Number of gaps | - | 130 | 122 |

**Table S2**. BUSCO scores for genome assembly.

| BUSCO | HAP1_Assembly | HAP1_Annotation | HAP2_Assembly | HAP2_Annotation |
|---|---|---|---|---|
| Complete BUSCOs (C) | 1586 (98.27%) | 1570 (97.27%) | 1599 (99.07%) | 1585 (98.2%) |
| Complete and single-copy BUSCOs (S) | 1544 (95.66%) | 1536 (95.17%) | 1543 (95.6%) | 1543 (95.6%) |
| Complete and duplicated BUSCOs (D) | 42 (2.6%) | 34 (2.11%) | 56 (3.47%) | 42 (2.6%) |
| Fragmented BUSCOs (F) | 11 (0.68%) | 10 (0.62%) | 8 (0.5%) | 10 (0.62%) |
| Missing BUSCOs (M) | 17 (1.05%) | 34 (2.11%) | 7 (0.43%) | 19 (1.18%) |

**Table S3**. Density information of repeats/transposable elements (haplotype 1)

| TE Classification | Coverage (bp) | Copy Number | % Genome Coverage | Genome Size | TE Family Count |
|---|---|---|---|---|---|
| DNA | 74310682 | 82109 | 6.72032 | 1105761065 | 82108 |
| Rolling Circle | 2638159 | 9224 | 0.238583098 | 1105761065 | 9224 |
| Penelope | 12038202 | 40669 | 1.08868022 | 1105761065 | 40669 |
| LINE | 38377514 | 61276 | 3.470687766 | 1105761065 | 61276 |
| SINE | 577135 | 4802 | 0.052193464 | 1105761065 | 4802 |
| LTR | 463373470 | 283009 | 41.90538848 | 1105761065 | 283009 |
| Other (Simple Repeat, Microsatellite, RNA) | 45597076 | 240884 | 4.123592107 | 1105761065 | 240884 |
| Unclassified | 179066643 | 521797 | 16.19397252 | 1105761065 | 521797 |
| Non-Repeat | 289782184 | NA | 26.20658234 | 1105761065 | NA |

**Table S4**. Species information used for phylogenetic analysis

| Species | Database or Download DOI (Assembly) | Publication |
|---|---|---|
| *Astilbe chinensis* | https://doi.org/10.6084/m9.figshare.28748501 | https://doi.org/10.1038/s41467-025-64842-9 |
| *Bergenia purpurascens* | https://doi.org/10.6084/m9.figshare.27092434 | https://doi.org/10.1038/s41597-025-04473-3 |
| *Bergenia scopulosa* | https://doi.org/10.5524/102589 | https://doi.org/10.1093/gigascience/giae091 |
| *Chrysosplenium sinicum* | https://doi.org/10.6084/m9.figshare.24558607.v2 | https://doi.org/10.1038/s42003-024-06701-9 |
| *Coffea canephora* | https://plants.ensembl.org/ (release 62) (AUK_PRJEB4211_v1) | - |
| *Glycine max* | https://plants.ensembl.org/ (release 62) (Glycine_max_v2.1) | https://doi.org/10.1038/nature08670 |
| *Hamamelis virginiana* | https://doi.org/10.6084/m9.figshare.13502001 | https://doi.org/10.1016/j.cub.2021.01.104 |
| *Kalanchoë fedtschenkoi* | http://phytozome.jgi.doe.gov/ (v1.1) | https://doi.org/10.1038/s41467-017-01491-7 |
| *Kalanchoë laxiflora* | http://phytozome.jgi.doe.gov/ (v1.1) | - |
| *Liquidambar styraciflua* | https://doi.org/10.6084/m9.figshare.26076514 | https://doi.org/10.1038/s41597-024-03924-7 |
| *Myriophyllum alterniflorum* | https://portal.darwintreeoflife.org/data (GCA_964291835.1) | - |
| *Paeonia ludlowii* | https://doi.org/10.6084/m9.figshare.23537670 | https://doi.org/10.1093/hr/uhad241 |
| *Paeonia ostii* | https://ftp.cngb.org/pub/CNSA/data5/CNP0003098/CNS0560369/CNA0050666/ | https://doi.org/10.1038/s41467-022-35063-1 |
| *Parrotia persica* | https://ngdc.cncb.ac.cn (GWHGEPQ00000000.1) | https://doi.org/10.1111/tpj.70367 |
| *Parrotia subaequalis* | https://ngdc.cncb.ac.cn (GWHGEPP00000000.1) | https://doi.org/10.1111/tpj.70367 |
| *Rhodiola kirilowii* | https://ngdc.cncb.ac.cn (GWHFGNY00000000.1) | https://doi.org/10.1038/s41597-025-04962-5 |
| *Ribes nigrum* | https://git.nfdi4plants.org/usadellab/ribes_nigrum_genome | https://doi.org/10.1093/hr/uhae313 |
| *Santalum album* | https://doi.org/10.6084/m9.figshare.23694729.v1 | https://doi.org/10.1038/s41597-023-02849-x |
| *Solanum lycopersicum* | https://plants.ensembl.org/ (release 62) (SL3.0) | - |
| *Theobroma cacao* | https://plants.ensembl.org/ (release 62) (Theobroma_cacao_20110822) | https://doi.org/10.1186/gb-2013-14-6-r53; https://doi.org/10.1038/ng.736 |
| *Tiarella polyphylla* | https://ngdc.cncb.ac.cn (GWHBKAH00000000) | https://doi.org/10.1111/1755-0998.13833 |
| *Umbilicus rupestris* | https://portal.darwintreeoflife.org/data (drUmbRupe1) | - |
| *Vitis vinifera* | https://plants.ensembl.org/ (release 62) (ASM3070453v1) | https://doi.org/10.1093/hr/uhad061 |
| *Penthorum chinense* | requested from authors | https://doi.org/10.1093/hr/uhad274 |

**Table S5.** Samples used for sequencing in this study.

| Sequencing code | Plant code | Tissue | Type | Sample name | SRA accession | Use |
|---|---|---|---|---|---|---|
| Bark_1 | B3 | Bark | cDNA (polyA) | B3Ba1 | SAMN58575598 | Genome annotation |
| Bark_2 | B3 | Bark | cDNA (polyA) | B3Ba2 | SAMN58575599 | Genome annotation |
| Bark_3 | B3 | Bark | cDNA (polyA) | B3Ba3 | SAMN58575600 | Genome annotation |
| Bud_1 | B3 | Bud | cDNA (polyA) | B3Bu1 | SAMN58575601 | Genome annotation |
| Bud_2 | B3 | Bud | cDNA (polyA) | B3Bu2 | SAMN58575602 | Genome annotation |
| Bud_3 | B3 | Bud | cDNA (polyA) | B3Bu3 | SAMN58575603 | Genome annotation |
| ILeaf_1 | B3 | immature leaf | cDNA (polyA) | B3IL1 | SAMN58575604 | Genome annotation |
| ILeaf_2 | B3 | immature leaf | cDNA (polyA) | B3IL2 | SAMN58575605 | Genome annotation |
| ILeaf_3 | B3 | immature leaf | cDNA (polyA) | B3IL3 | SAMN58575606 | Genome annotation |
| MLeaf_1 | B3 | mature leaf | cDNA (polyA) | B3ML1 | SAMN58575607 | Genome annotation |
| MLeaf_2 | B3 | mature leaf | cDNA (polyA) | B3ML2 | SAMN58575608 | Genome annotation |
| MLeaf_3 | B3 | mature leaf | cDNA (polyA) | B3ML3 | SAMN58575609 | Genome annotation |
| Petiole_1 | B3 | Petiole | cDNA (polyA) | B3P1 | SAMN58575610 | Genome annotation |
| Petiole_2 | B3 | Petiole | cDNA (polyA) | B3P2 | SAMN58575611 | Genome annotation |
| Petiole_3 | B3 | Petiole | cDNA (polyA) | B3P3 | SAMN58575612 | Genome annotation |
| Stem_1 | B3 | Stem | cDNA (polyA) | B3S1 | SAMN58575613 | Genome annotation |
| Stem_2 | B3 | Stem | cDNA (polyA) | B3S2 | SAMN58575614 | Genome annotation |
| Stem_3 | B3 | Stem | cDNA (polyA) | B3S3 | SAMN58575615 | Genome annotation |
| WStem_1 | B3 | Woody Stem | cDNA (polyA) | B3WS1 | SAMN58575616 | Genome annotation |
| WStem_2 | B3 | Woody Stem | cDNA (polyA) | B3WS2 | SAMN58575617 | Genome annotation |
| WStem_3 | B3 | Woody Stem | cDNA (polyA) | B3WS3 | SAMN58575618 | Genome annotation |
| Root_1 | B3 | Root | cDNA (polyA) | B3R1 | SAMN58575619 | Genome annotation |
| Root_2 | B3 | Root | cDNA (polyA) | B3R2 | SAMN58575620 | Genome annotation |
| Root_3 | B3 | Root | cDNA (polyA) | B3R3 | SAMN58575621 | Genome annotation |
| HMW_DNA | B3 | Leaf | HMW-DNA | B3IL-HMW | SAMN58575622 | Genome assembly |
| HIC_DNA | B3 | Leaf | DNA for HiC | B3ML-HIC | SAMN58575623 | Genome assembly |
| S1 | Fa | Bark | cDNA (polyA) | Fa_B | SAMN58575580 | Paired omics |
| S2 | Fa | immature leaf | cDNA (polyA) | Fa_IM | SAMN58575581 | Paired omics |
| S3 | Fa | mature leaf | cDNA (polyA) | Fa_ML | SAMN58575582 | Paired omics |
| S4 | Fa | petiole | cDNA (polyA) | Fa_P | SAMN58575583 | Paired omics |
| S5 | Fa | Root | cDNA (polyA) | Fa_R | SAMN58575584 | Paired omics |
| S6 | Fa | stem | cDNA (polyA) | Fa_S | SAMN58575585 | Paired omics |
| S7 | Fb | Bark | cDNA (polyA) | Fb_B | SAMN58575586 | Paired omics |
| S8 | Fb | immature leaf | cDNA (polyA) | Fb_IM | SAMN58575587 | Paired omics |
| S9 | Fb | mature leaf | cDNA (polyA) | Fb_ML | SAMN58575588 | Paired omics |
| S10 | Fb | petiole | cDNA (polyA) | Fb_P | SAMN58575589 | Paired omics |
| S11 | Fb | Root | cDNA (polyA) | Fb_R | SAMN58575590 | Paired omics |
| S12 | Fb | stem | cDNA (polyA) | Fb_S | SAMN58575591 | Paired omics |
| S13 | Fc | Bark | cDNA (polyA) | Fc_B | SAMN58575592 | Paired omics |
| S14 | Fc | immature leaf | cDNA (polyA) | Fc_IM | SAMN58575593 | Paired omics |
| S15 | Fc | mature leaf | cDNA (polyA) | Fc_ML | SAMN58575594 | Paired omics |
| S16 | Fc | petiole | cDNA (polyA) | Fc_P | SAMN58575595 | Paired omics |
| S17 | Fc | Root | cDNA (polyA) | Fc_R | SAMN58575596 | Paired omics |
| S18 | Fc | stem | cDNA (polyA) | Fc_S | SAMN58575597 | Paired omics |
